# Maximum Likelihood Estimation for Unrooted 3-Leaf Trees: An Analytic Solution for the CFN Model

**DOI:** 10.1101/2024.02.01.578504

**Authors:** Max Hill, Sebastien Roch, Jose Israel Rodriguez

## Abstract

Maximum likelihood estimation is among the most widely-used methods for inferring phylogenetic trees from sequence data. This paper solves the problem of computing solutions to the maximum likelihood problem for 3-leaf trees under the 2-state symmetric mutation model (CFN model). Our main result is a closed-form solution to the maximum likelihood problem for unrooted 3-leaf trees, given generic data; this result characterizes all of the ways that a maximum likelihood estimate can fail to exist for generic data and provides theoretical validation for predictions made in [28]. Our proof makes use of both classical tools for studying group-based phylogenetic models such as Hadamard conjugation and reparameterization in terms of Fourier coordinates, as well as more recent results concerning the semi-algebraic constraints of the CFN model. To be able to put these into practice, we also give a complete characterization to test genericity.

## 1. Introduction and preliminaries

This paper is concerned with inferring the evolutionary history of a set of a species or other taxa from sequence data using maximum likelihood. In practice, maximum likelihood inference is among the most commonly-used in phylogenetic analyses, and in contrast to the simple (but more analytically tractable) model considered in this paper, maximum likelihood estimation is typically undertaken with sophisticated models of site evolution, utilizing heuristic (e.g. hill-climbing) methods and multiple start points to explore the tree space in order to obtain parameters which maximize the likelihood. A variety of excellent and widely-used implementations exist, many of which have been used in thousands of studies (e.g., [27, 32, 39, 29]).

Nonetheless, there remains interest in computing *analytic* (i.e., closed-form) solutions in simpler cases [11, 10, 8, 37, 7, 20, 9] as well as exact solutions via algebraic methods [23, 15], with the goal of providing a more rigorous understanding of the properties of maximum likelihood estimation and the ways that it can fail. For example, it is well-known that the maximum likelihood tree need not be unique [33], that for certain data there exists a continuum of trees which maximize the likelihood [8], and that there exist data for which the maximum likelihood estimate does not exist—or at least, is not a true tree with finite branch lengths [23]. In the context of phylogenetic estimation, maximum likelihood exhibits complex behavior even in very simple cases; one important example of this is long-branch attraction, a form of estimation bias which has been the subject of much interest (see, e.g., [36, 28, 5, 3]), but is not yet fully understood.

The specific problem that we consider in this paper is that of using maximum likelihood to estimate the branch lengths of an unrooted 3-leaf tree from molecular data generated according to the Cavendar-Farris-Neyman (CFN) model, a binary symmetric model of site substitution. A related problem of estimating rooted 3-leaf trees under the molecular clock assumption was considered by [37]. The problem considered here differs from that in [37] as we do not assume a molecular clock; instead our problem involves maximizing the likelihood over three independently varying branch length parameters.

The 3-leaf MLE problem in this paper was considered, though not fully solved, in [23]. There, the authors introduced a general algorithm for computing numerically exact solutions using semi-algebraic constraints (i.e. polynomial inequalities) satisfied by phylogenetic models, along with methods from numerical algebraic geometry. Applying their method to the 3-leaf maximum likelihood problem under the CFN model, they discovered a nontrivial example where the maximum likelihood estimate does not exist. A similar problem was also considered in [28], who obtained a partial solution (i.e., involving simulations) for the 4-state Jukes-Cantor model, allowing them to use distance estimates to predict with high accuracy certain features of the maximum likelihood estimate for specific data.

In this paper, we go one step further, presenting a full solution to the 3-leaf maximum likelihood problem under the CFN model, for generic data. Our proofs have a similar flavor to the model boundary decompositions technique in [1, Section 4], but we consider a submodel that is not full dimensional. The main result of this paper provides a full characterization of the behavior of maximum likelihood estimation in this setting, up to and including necessary and sufficient conditions for the MLE to exist, as well as detailed analysis describing the ways in which an MLE may fail to exist. Further, our results validate the predictions given in [28], providing theoretical underpinning to an interesting connection between maximum likelihood and distance-based estimates appearing in that paper.

While finalizing this manuscript, the recent work of [20] came to our attention. Building on a connection between the multinomial distribution and maximum likelihood estimation of 3-leaf trees under the CFN model, the authors of [20] make a surprising and interesting discovery about the likelihood geometry for 3-leaf models: in the case of three leaves, estimation by maximum likelihood is equivalent to estimation by pairwise distances, an equivalence which does not hold for trees with four or more leaves. In particular, the authors use this to obtain an analytic solution to the MLE problem for 3-leaf trees which is applicable whenever the data (regarded here as a vector representing the empirical site pattern frequencies) lies in the interior of the parameter space of the model. The present paper provides a natural extension of the 3-leaf result in [20] in two ways: first, by providing a simple characterization of when the data lies in the interior of the model, and second, by analyzing in detail the case when it does not; the analysis of this non-interior case is both substantial and technically non-trivial, and provides an improved understanding of the settings where maximum likelihood is prone to failure, such as in the case of trees with very long or short branches.

The remainder of this paper is structured as follows. In Section 1 we introduce the model, problem statement, and some of the key tools that will be employed. In Section 2, we present our main result, a closed-form “analytic” solution to the maximum-likelihood problem for 3-leaf trees under the CFN model. In Section 3, we discuss the significance and novel contribution of this result. A proof of the main result is then presented in Section 4.

### 1.1. Data and model of evolution

#### 1.1.1. Tree parameter

For any finite set 𝒳, an 𝒳*-tree* 𝒯 is an ordered pair (*T* ; *φ*) where *T* is a tree with vertex set *V* (*T*) and the *labelling map φ* : 𝒳 → *V* is a map such that *v* ∈ *φ*(𝒳) whenever *v* ∈ *V* (*T*) and deg(*v*) ≤ 2. If *φ* is a bijection into the leaves of *T*, then 𝒯 is called a *phylogenetic* 𝒳*-tree*; in this case the elements of 𝒳 are identified with the leaves of the tree (for a standard reference, see [31]). In addition, we associate with 𝒯 a vector of nonnegative *branch lengths d* := (*d*_*e*_)_*e*∈*E*(*T*)_, where *E*(*T*) is the edge set of *T*.

We regard 𝒳 as a set of taxa, with *T* representing a hypothesis about their evolutionary or genealogical history; the branch lengths are regarded as representing a measure of evolutionary distance measured in *expected number of mutations per site*. In this paper we consider exclusively the case 𝒳 = [3].

Rather than using the evolutionary distances *d* as edge parameters of *T*, for our analyses it will be more convenient to use an alternative parameterization of the branch lengths as a vector *θ* := (*θ*_*e*_)_*e*∈*E*(*T*)_ ∈ [0, 1]^|*E*(*T*)|^, where

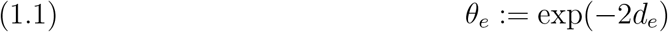

for all *e* ∈ *E*(*T*). The numerical edge parameters (*θ*_*e*_)_*e*∈*E*(*T*)_ have been referred to as “path-set variables” [10]; we refer to them here as the *Hadamard parameters*.

#### 1.1.2. Site substitution model

The site substitution model considered in this paper is the fully symmetric *Cavendar-Farris-Neyman (CFN)* model. Also known as the *N*_2_ model (e.g., in [31]), the CFN model takes as input a tree parameter 𝒯_*θ*_ = (𝒯, *θ*), consisting of a phylogenetic [*n*]-tree 𝒯= (*T, φ*) along with edge parameters *θ* = (*θ*_*e*_)_*e*∈*E*(*T*)_, and outputs a random vector *X* = (*X*_1_, …, *X*_*n*_), whose entries *X*_1_, …, *X*_*n*_ ∈ {−1, +1} are associated with the *n* leaves of 𝒯.

The CFN model, corresponding to a time-reversible Markov chain on a tree, is the simplest model of site substitution, possessing only two nucleotide states, which we denote by +1 (pyrimidine) and −1 (purine). Under this model, the probability of a nucleotide in state *i* transitioning to state *j* over an edge of length *t* can be shown to be

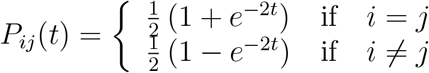

for all *i, j* ∈ {−1, +1} ([37], and for a more general reference, see [31, p.197]). In other words, the transition probability from state *i* to *j* along a given edge *e*, denoted *P*_*ij*_(*e*), is

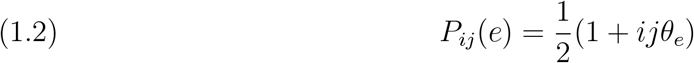

for all *i, j* ∈ {+1, −1}.

Moreover, we assume a uniform root distribution, from which it follows that

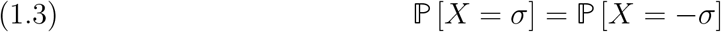

for all *σ* ∈ {−1, 1}^*n*^ (c.f. Lemma 8.6.1.(ii) in [31], see also [34, p.221]).

The distribution of *X* depends on both the topology and branch lengths of the tree parameter. For a phylogenetic [*n*]-tree 𝒯 with topology *τ* and Hadamard parameters *θ*, we use the notation

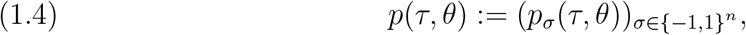

where

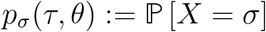

and ℙ is the distribution of *X* under the CFN model with parameters 𝒯 and *θ*.

#### 1.1.3. Identifiability of Model Parameters

Another way to understand the CFN model is as a parameterized statistical model with the tree topology *τ* held fixed; in this case, the CFN model is regarded as the image of the map

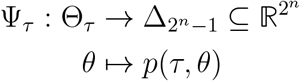

where Θ_*τ*_ ⊆ [0, 1]^|*E*(*T*)|^ is the set of possible Hadamard parameters, and where Δ_*r*−1_ ⊂ ℝ^*r*^ is the probability simplex of dimension *r* − 1; i.e.,

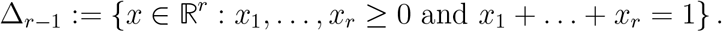

The usual assumption prescribed for the CFN model (which is *not* made in this paper) is that Θ_*τ*_ = (0, 1) ^|*E*(*T*)|^ (so that 0 *< θ*_*e*_ *<* 1 for all *e* ∈ *E*(*T*), or equivalently, that 0 *< d*_*e*_ *<* ∞ for all *e* ∈ *E*(*T*)). Under that assumption, Ψ_*τ*_ is injective [17], and hence the edge parameters *θ* = (*θ*_*e*_)_*e*∈*E*(*T*)_ are *identifiable*. This means that if 𝒯_*θ*_ := (𝒯, *θ*) and 𝒯_*θ′*_ := (𝒯, *θ*^*′*^) are two *n*-leaf trees with the same topology *τ* but with edge parameters *θ* and *θ*^*′*^ such that *θ* ≠ *θ*^*′*^, then the distributions of *X* will be different under 𝒯_*θ*_ and 𝒯_*θ′*_.

On the other hand in this paper, we consider an extension by allowing *θ*_*e*_ ∈ [0, 1] for each edge *e* ∈ *E*(*T*), rather than *θ*_*e*_ ∈ (0, 1). This extension allows for branch lengths which are infinite or zero (when measured in expected number of mutations per site), in order to better understand the behavior of maximum likelihood estimation in the limit as one or more branch lengths tend to zero or to infinity. This seemingly slight extension of the model substantially adds to the complexity of the analysis. In particular, as a consequence of this extension, it is no longer the case that the numerical parameters *θ* are identifiable, which presents certain complications, described in detail later in the paper. On the other hand, it also has the effect of guaranteeing the existence of the maximum likelihood estimate.

#### 1.1.4. Data

In practice, DNA sequence data is typically arranged as a *multiple sequence alignment*, an *n* × *N* matrix, with each row corresponding to a leaf of 𝒯 and each column representing an aligned site position. It is standard (albeit unrealistic) to assume that sites evolved independently, and as such our data consists of *N* random column vectors

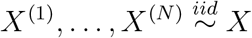

where *X* is a random variable taking values in {+1, −1}^*n*^ whose distribution will be described below, and which is regarded as a vector of nucleotides observed at the leaves of 𝒯 such that *X*_*i*_ is the nucleotide observed at the vertex with label *i* for each *i* ∈ [*n*]. Under the CFN model, the distribution of *X* depends on the topology of as well as the edge parameters *θ*. Due to the exchangeability of *X*^(1)^, …, *X*^(*N*)^, the data can be summarized by a *site frequency vector*

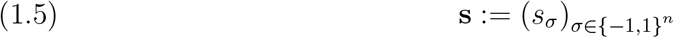

where

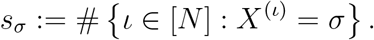

#### 1.1.5. α-Split Patterns

In light of Eq. (1.3), it is possible using a change of coordinates to represent the distribution of *X* using a vector of 2^*n*−1^ entries rather than 2^*n*^ entries. For any vector *σ* ∈ {−1, +1}^*n*^ and any *α* ∈ [*n* −1], we say that *σ* has *α-split pattern* if there exists *k* ∈ {−1, +1} such that *σ*_*i*_ = *k* if and only if *i* ∈ *α*.

For example, if *n* = 3 then (+1, −1, +1) and (−1, +1, −1) both have {2} -split pattern; (+1, +1, −1) and (−1, −1, +1) both have {1, 2} -split pattern, and so forth.

Analogously to the notation in Eq. (1.4), define the *expected site pattern spectrum* to be

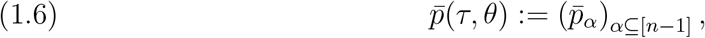

where

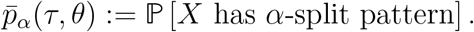

In other words, 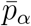 is the probability that the entries of *X* correspond to the split *α*|([*n*]\*α*). For this vector 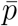, we follow [18] in assuming that the subsets of [*n* − 1] are ordered lexicographically, e.g., so that for *n* = 3, we have the order (∅, {1*}, {*2*}, {*1, 2*}*).

By Eq. (1.3), the distribution vector 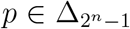 is fully specified without loss of information by the lower dimensional vector 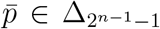 (c.f., [2]). The data may also be summarized by the lower-dimensional (but sufficient) statistic

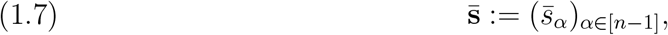

where

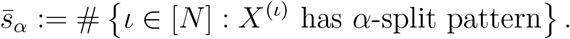

### 1.2. The maximum likelihood problem

Let 𝒯 = (*T, φ*) be a phylogenetic [*n*]-tree with unrooted tree topology *τ* and Hadamard edge parameters *θ* = (*θ*_*e*_)_*e*∈*E*(*T*)_ for some *n* ≥ 2. Given data taking the form of Eq. (1.5), or equivalently Eq. (1.7), the *log-likelihood function* is the function

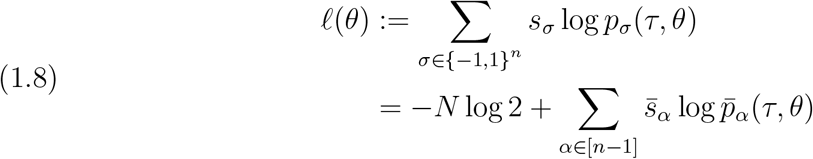

where *p*_*σ*_ and 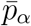 are defined as in Eqs. (1.4) and (1.6).

The *maximum likelihood problem* is to find all parameters *τ* and *θ* ∈ [0, 1]^|*E*(*T*)|^ which maximize Eq. (1.8).

When *n* = 3, there is only one unrooted tree topology, in which case this problem reduces to that of finding the numerical parameters *θ* ∈ [0, 1]^3^ which maximize Eq. (1.8).

### 1.3. Hadamard conjugation

In this section we introduce an important reparametrizaton of the CFN model, as well as a central tool in our analyses: Hadamard conjugation. For any even subset *Y* ⊆ [*n*], define the *path set P* (𝒯, *Y*) induced by *Y* on 𝒯 to be the set of 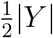 edge-disjoint paths in 𝒯, each of which connects a pair of leaves labelled by elements from *Y*, taking *P* (𝒯, ∅) = ∅. This set is unique if 𝒯 is a binary tree [31].

The *edge spectrum* is the vector

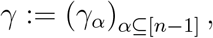

where

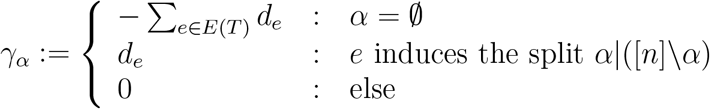

Inductively define *H*_0_ := [1], and for *k* ≥ 0 define

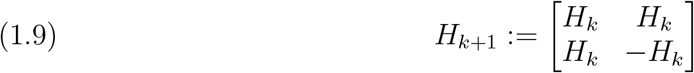

Let *H* := *H*_*n*−1_. Then *H* is a 2^*n*−1^ × 2^*n*−1^ matrix, and by our choice of ordering for [*n* − 1], we have *H* = (*h*_*α,β*_)_*α,β*⊆[*n*−1]_ where

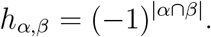

In particular, *H* is a symmetric Hadamard matrix with 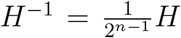. Since *H* is the character table of the group 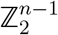, multiplication by *H* can be regarded as a discrete Fourier transformation, a commonly-used tool in the study of the CFN model and other group-based substitution models (see [31, chp. 8] as well as [12, 35, 19]). Closely related to the discrete Fourier transform is the next theorem, which allows us to translate between the edge spectrum and the expected site pattern spectrum.

#### Theorem 1.1

(Hadamard conjugation [18, 14]). Let 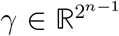 be the edge spectrum of a phylogenetic [*n*]-tree 𝒯, let *H* := *H*_*n*−1_, and let 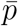 the expected site pattern spectrum as defined in Eq. (1.6). Then

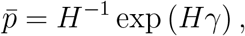

where the exponential function exp(·) is applied component-wise.

Hadamard conjugation, as formulated in this theorem, has proved to be an essential tool in a number of previous results and has analogues for substitution models other than the CFN (see, e.g., [10, 8]). For a proof and a detailed discussion of this theorem, we refer the reader to [31]. In particular, we will utilize the following proposition, itself a consequence of Theorem 1.1.

#### Proposition 1.2

(Corollary 8.6.6 in [31]). Let *θ*_*e*_ ∈ [0, 1] for all *e* ∈ *E*(*T*). Then for all subsets *α* of [*n* − 1],

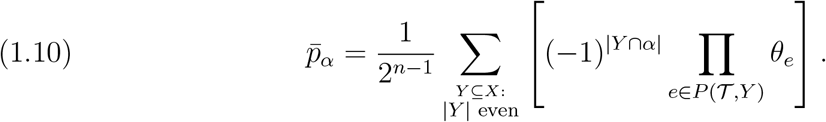

Note that Proposition 1.2 holds even if the root distribution is not taken to be uniform, however we do not consider that case in this paper. When *n* = 3, Eq. (1.10) reduces to

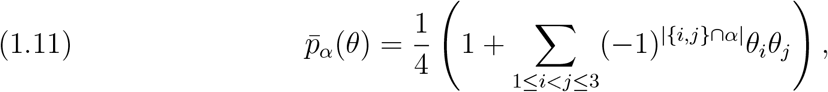

and by a change of notation, this can be rewritten as

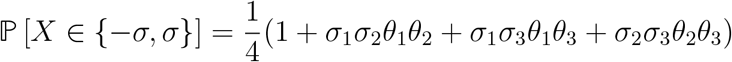

for all *σ* ∈ {−1, +1}^3^. Therefore by Eq. (1.3),

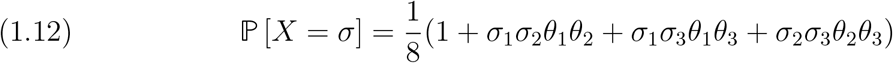

for all *σ* ∈ {+1, −1}^3^.

### 1.4. Interpretation of Hadamard parameters

We regard a vector of Hadamard parameters *θ* = (*θ*_*e*_)_*e*∈*E*(*T*)_ as *biologically plausible* if *θ*_*e*_ ∈ (0, 1) for all *e* ∈ *E*(*T*). Since 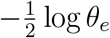 is the expected number of mutations on edge *e*, it follows that *θ*_*e*_ ∈ (0, 1) if and only if *d*_*e*_ ∈ (0, *∞*). In other words, biologically plausible Hadamard parameters correspond to trees with branch lengths having positive and finite expected number of mutations per site. In this work, we allow for *θ*_*e*_ ∈ [0, 1] in order to better study the ways that maximum likelihood can fail to return a tree with biologically plausible parameters; for example, trees with extremely short or long branches are of special interest, since it is in this setting that long-branch attraction is hypothesized to occur.

Observe that *θ*_*e*_ ∈ [0, 1] measures the correlation between the state of the Markov process at the endpoints of the edge *e*. Suppose *e* = (*u, v*) ∈ *E*(*T*) and let *X*_*u*_ and *X*_*v*_ denote the state of the Markov process at nodes *u* and *v* respectively. Eq. (1.2) implies that if *θ*_*e*_ = 1 then *X*_*u*_ = *X*_*v*_ with probability 1. On the other hand, if *θ*_*e*_ = 0 then *X*_*u*_ and *X*_*v*_ are independent; to see why this is the case, observe that using Eq. (1.2), it holds for all *i, j* ∈ {−1, 1} that

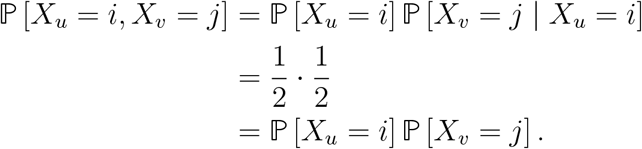

The observation has an important consequence which is summarized in the next lemma. In particular, by conditioning on the state of the Markov process at the endpoints of *e* and using the Markov property, a straightforward calculation gives the following result:

#### Lemma 1.3

(Independence caused by “infinitely long” branches). Let 𝒯 = (*T, φ*) be a phylogenetic [*n*]-tree. Let *e* ∈ *E*(*T*) and let *A*_*e*_|*B*_*e*_ denote the split induced by *e* on the leaf set [*n*]. If *θ*_*e*_ = 0 then the random vectors (*X*_*i*_ : *i* ∈ *A*_*e*_) and (*X*_*i*_ : *i* ∈ *B*_*e*_) are independent.

A proof of Lemma 1.3 can be found in Appendix A.

### 1.5. Fourier coordinates and semi-algebraic constraints

One key takeaway of 1.2 is that the probability of any site pattern can be computed as a *polynomial* function of the Hadamard parameters.

Hence the CFN model for a fixed tree topology *τ* may be regarded as the image of the map

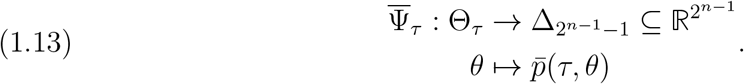

In our setting, we take Θ_*τ*_ = [0, 1]^|*E*(*T*)|^ and regard the statistical model as the image

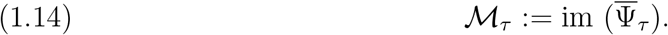

An important monomial parameterization of the CFN model is obtained by means of the discrete Fourier representation (see [12, 35, 34]), which here is given by the matrix *H*. If **q** = (*q*_111_, *q*_101_, *q*_011_, *q*_110_)^⊤^ is a vector of *Fourier coordinates*, then 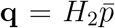. In the case with *n* = 3, we have *γ* = (−(*d*_1_ + *d*_2_ + *d*_3_), *d*_1_, *d*_2_, *d*_3_)^⊤^, so that Theorem 1.1 implies that

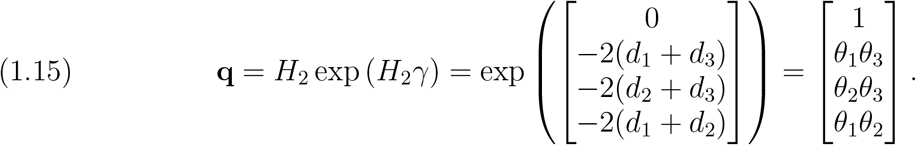

For group-based models (see, e.g., [35]) like the CFN model, 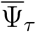 is a polynomial map, and hence all points in ℳ_*τ*_ satisfy certain polynomial equalities, called *phylogenetic invariants*, and polynomial inequalities, called *semi-algebraic constraints*, both of which are usually formulated in terms of Fourier coordinates.

In the 3-leaf case considered here, the only phylogenetic invariant is *q*_111_ = 1, which is equivalent to the stochastic invariant 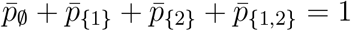.

The semi-algebraic constraints of the CFN model were studied more generally by [25, 23]. When the assumption is made that the tree parameter has biologically plausible parameters, the semi-algebraic constraints for 3-leaf trees consist of the following inequalities:

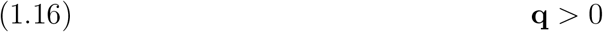

and

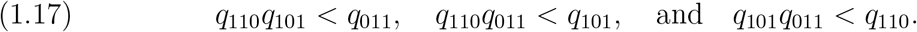

The inequality Eq. (1.16) corresponds simply to the assumption that the evolutionary distance between each pair of leaves is finite.

The remaining semi-algebraic constraints in Eq. (1.17) have straightforward interpretation in terms of the additive evolutionary distances on the tree induced by the branch lengths *d*_1_, *d*_2_, and *d*_3_. To see this, observe that by Eq. (1.15), the inequalities in Eq. (1.17) can be written as

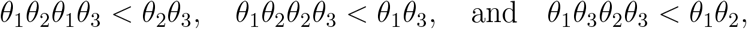

and by Eq. (1.1), these are equivalent to

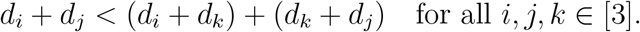

In other words, the semi-algebraic constraints in Eq. (1.17) are nothing but the triangle inequality in disguise.^1^

Taken together, Eqs. (1.16) and (1.17) are equivalent to the tree having branch lengths in *d*_1_, *d*_2_, *d*_3_ which are both positive and finite. Note this assumption is not made in this paper, as we allow for branch lengths to also be either zero or infinite, a relaxation which corresponds to using non-strict inequalities in Eqs. (1.16) and (1.17). Nonetheless, the strict inequalities will play an important role in our main result.

#### Remark 1.4.

The semi-algebraic constraints in (1.17) show up in other settings as well. For example, in [25] they appear as embeddability conditions for the Kimura 3-parameter model; in that setting, the inequalities are equivalent to the nonnegativity of the off-diagonal entries of the mutation rate matrix, and therefore—just as in our setting— implicitly specify that the branch lengths be nonnegative. For a generalization of these inequalities as embeddability conditions see the main result of [4].

Because the Fourier coordinates factorize into Hadamard parameters, as shown in Eq. (1.15) (and more generally: see, e.g., [31]), the nontrivial Fourier coordinates thus have a simple biological interpretation. For each distinct pair *i, j* ∈ [3], since 𝔼 [*X*_*i*_] = 𝔼 [*X*_*j*_] = 0, it follows that

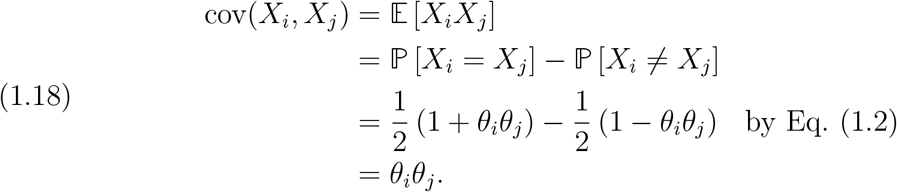

In other words, the nontrivial Fourier coordinates are covariances of nucleotides observed at the leaves of the tree.

Both the monomial parameterization in Eq. (1.15) and the semialgebraic constraints in Eqs. (1.16) and (1.17) will play an important role in the proof and interpretation of our main result for 3-leaf trees. In particular, our approach is to estimate the Fourier coordinates *θ*_1_*θ*_2_, *θ*_1_, *θ*_3_, *θ*_2_*θ*_3_ directly from the data, and it turns out that whether or not these estimates satisfy inequalities corresponding to Eqs. (1.16) and (1.17) completely determines the qualitative properties of the maximum likelihood estimate.

## 2. Main result: an analytic solution to the 3-leaf MLE problem

### 2.1. The 3-leaf maximum likelihood problem

Let 𝒯 be an unrooted phylogenetic [3]-tree, with unknown numerical edge parameters 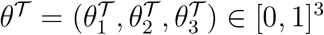, as shown in Fig. 1. Let **s** be a site frequency vector obtained from *N* independent samples *X*^(1)^, …, *X*^(*N*)^ generated according to the CFN process on 𝒯. The *3-leaf maximum likelihood problem* is to find all numerical parameters *θ* ∈ [0, 1]^3^ which maximize Eq. (1.8) given the data **s**.

**Figure 1.**
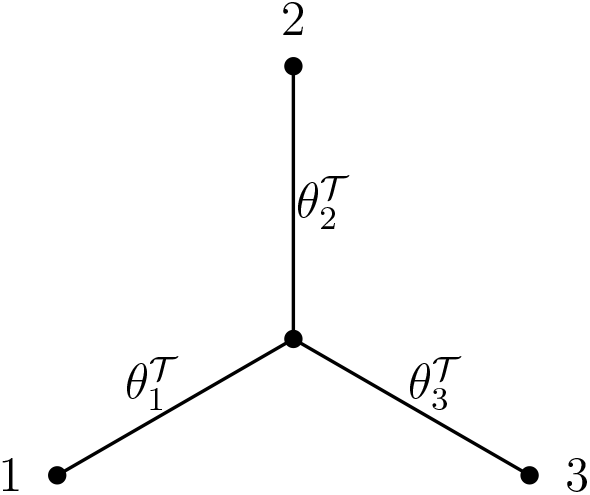
Three-leaf tree 𝒯 with Hadamard edge parameters 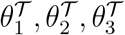.

Note that since there is only one possible unrooted topology for a 3-leaf tree, the topology parameter *τ* does not play a role in this problem.

### 2.2. Key definitions and notation

The key statistics used are the following:

#### Definition 2.1

(The statistics 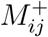,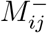,*B*_*ij*_, **B**). For all *i, j* ∈ [*n*] such that *i*≠*j*, define

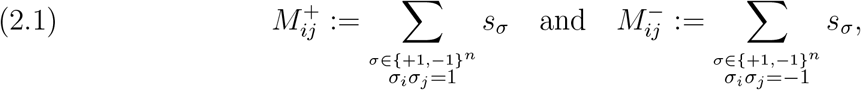

as well as

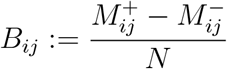

and

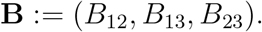

In words, 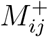 is the number of samples for which leaves *i* and *j* share the same nucleotide state and 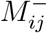 is the number for which the nucleotides observed at leaves *i* and *j* differ. It follows by definition that 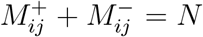 and that 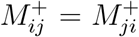 and 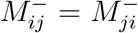 for all distinct *i, j* ∈ [3].

The statistic *B*_*ij*_ measures the observed correlation of the observations at leaves *i* and *j* of the tree. By the law of large numbers,

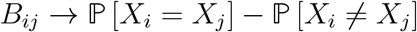

almost surely as *N* → ∞. Therefore by Eq. (1.18), *B*_*ij*_ is a consistent estimator of the Fourier coordinate *θ*_*i*_*θ*_*j*_, which is itself the covariance of *X*_*i*_ and *X*_*j*_. Moreover, it is easy to check that

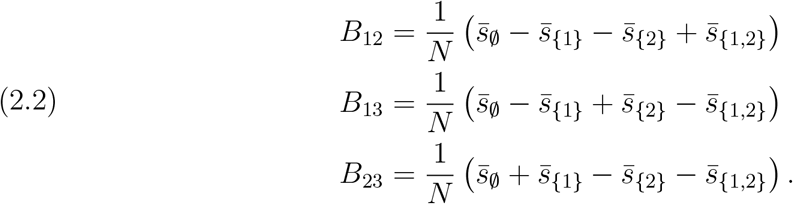

Due to symmetries of the problem, it will be useful to index the statistics in Definition 2.1 using permutations.

**Notation 2.2** (Indexing with permutations). Let Alt(3) denote the alternating group of degree 3, which can be expressed in cycle notation as

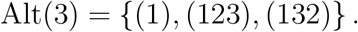

For each *π* ∈ Alt(3), we write

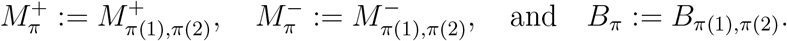

The use of the statistic **B** will permit us to obtain a simple criterion for when a maximum likelihood estimate corresponding to a 3-leaf tree with finite branch lengths exists. This criteria involves the following set:

#### Definition 2.3

(The set 𝒟). Define

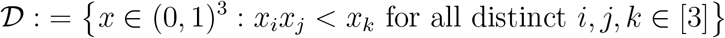

As we will show in the next theorem, it turns out that given some fixed data **s**, a maximum likelihood estimate with finite branch lengths exists precisely if and only if **B** ∈𝒟. Importantly, since **B** is an estimate of the nontrivial Fourier coordinates (*q*_110_, *q*_101_, *q*_011_) = (*θ*_1_*θ*_2_, *θ*_1_*θ*_3_, *θ*_2_*θ*_3_), the inequalities which define 𝒟 correspond precisely to the semi-algebraic constraints in Eqs. (1.16) and (1.17).

### 2.3. Assumptions about the data

We make two simplifying assumptions about the data **s**:

A.1 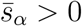 for all *α* ⊆ [2].
A.2 *B*_12_, *B*_13_ and *B*_23_ are nonzero and distinct.

In words, assumption **A.1** states that each site pattern *aaa, aab, aba, abb* is observed at least once in the data (where *a* and *b* represent different nucleotides). One consequence of this is that 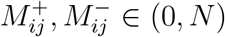 whenever *i ≠ j*. Since the number of site patterns is 2^*n−*1^, this assumption would be unrealistic for a tree with many more leaves (e.g., *n >* 30) given the size of genomic datasets [8], but for our purposes (i.e., with *n* = 3), this assumption is reasonable. Assumption **A.2** is an assumption about the genericity of the data, in the sense that it is equivalent to assuming that

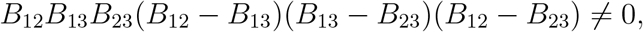

or equivalently,

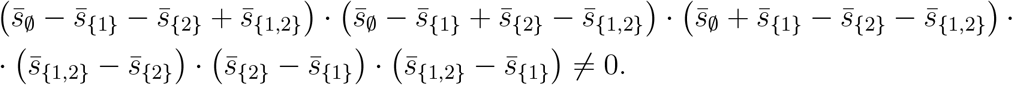

These two assumptions considerably simplify the problem with very little loss of generality, as both assumptions are likely to be satisfied when *N* is large.

### 2.4. Statement of main result

Our main result is the following theorem, which solves the maximum likelihood problem for 3-leaf trees; further discussion of this result is given in Section 3.

#### Theorem 2.4

(Global MLE for the 3-leaf tree). Assume that **A.1** and **A.2** hold. Then *ℓ* has a maximizer on the set [0, 1]^3^. Denote the set of all such maximizers as

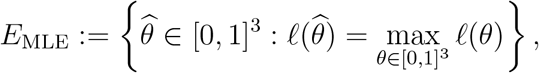

and let *π*_1_, *π*_2_, *π*_3_ ∈ Alt(3) such that

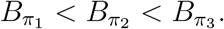

If **B** *∈*𝒟, then

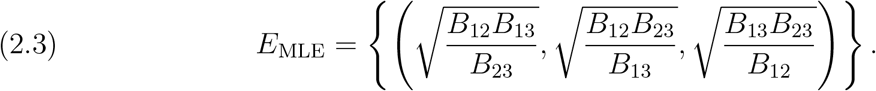

On the other hand, if **B** ∉ 𝒟, then the following trichotomy holds:

i. If 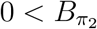 then

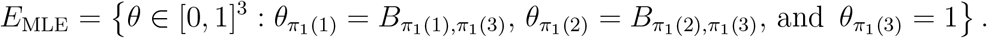
ii. If 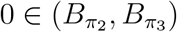 then

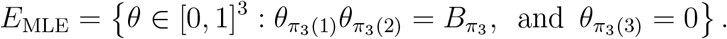
iii. If 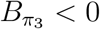 then

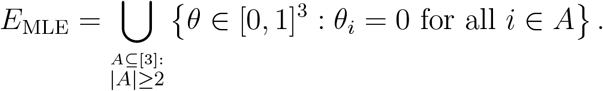

Enumerating out the possible cases of Theorem 2.4 immediately yields the following corollary.

#### Corollary 2.5.

Under the assumptions of Theorem 2.4, the MLE can be determined from the values of *B*_12_, *B*_13_, *B*_23_ using Table 1.

**Table 1.** Summary of results of Theorem 2.4, which gives the MLE for a 3-leaf tree, as a function of the data *B*_12_, *B*_13_, *B*_23_.

| Conditions | MLE $\theta_{\text{MLE}} = (\theta_1, \theta_2, \theta_3)$ |
| --- | --- |
| $B_{ij}B_{jk} < B_{ik}$ & $B_{ij} > 0$, all distinct $i, j, k \in [3]$ | $\left(\sqrt{\frac{B_{12}B_{13}}{B_{23}}}, \sqrt{\frac{B_{12}B_{23}}{B_{13}}}, \sqrt{\frac{B_{13}B_{23}}{B_{12}}}\right)$ |
| $B_{23} < B_{12}B_{13}$ & $B_{12}, B_{13} > 0$ | $(1, B_{12}, B_{13})$ |
| $B_{13} < B_{12}B_{23}$ & $B_{12}, B_{23} > 0$ | $(B_{12}, 1, B_{23})$ |
| $B_{12} < B_{13}B_{23}$ & $B_{13}, B_{23} > 0$ | $(B_{13}, B_{23}, 1)$ |
| $B_{12}, B_{13} < 0$ | $\theta_1 = 0, \theta_2\theta_3 = B_{23}$ |
| $B_{12}, B_{23} < 0$ | $\theta_2 = 0, \theta_1\theta_3 = B_{13}$ |
| $B_{13}, B_{23} < 0$ | $\theta_3 = 0, \theta_1\theta_2 = B_{12}$ |
| $B_{12}, B_{13}, B_{23} < 0$ | At least two of $\theta_1, \theta_2, \theta_3$ are zero,<br>with the third one taking any value |

Before proving this theorem in Section 4, we first discuss its significance and some implications.

## 3. Discussion of novel contribution

In addition to providing necessary and sufficient conditions for the MLE to exist as a tree with finite branch lengths, Theorem 2.4 also characterizes the ways that this can fail to occur, and highlights a subtle connection between the semi-algebraic constraints given in Eqs. (1.16) and (1.17) and properties of the maximum likelihood estimate.

In an important paper on long-branch attraction [28], a compelling connection was drawn between maximum likelihood and distance estimates on a 3-leaf tree under the Jukes-Cantor model of site substitution. Through a combination of analytic boundary case analysis and simulations, the authors argued that the failure of distance-based branch-length estimates to satisfy the triangle inequality and nonnegativity constraints was a good predictor of maximum likelihood failing to return a tree with biologically plausible branch lengths.

Due to the use of different substitution models (i.e., the CFN model considered here versus the 4-state Jukes-Cantor model considered in [28]), some translation is necessary to recognize the connection between our results and those of [28]. The distance estimates used in [28] are related to the standard *Jukes-Cantor correction* [22, 38], and are given by the formula

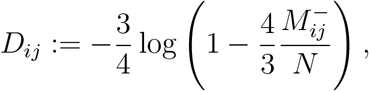

which returns an estimate of evolutionary distance *D*_*ij*_ between taxa *i* and *j*, measured in expected number of mutations per site. The variable 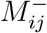 is the number of samples such that the nucleotide states observed at taxa *i* and *j* differ (i.e., the same as in this paper). The convention used in [28] is to define *D*_*ij*_ = *∞* if 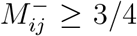. Formulated in terms of these distances, the predictions made in [28] are given in Table 2.

**Table 2.** Table of predictions (from [28]) for the behavior of maximum likelihood as a function of distance estimates.

| Conditions | Prediction |
| --- | --- |
| $D_{23} \geq D_{12} + D_{13}$ (incl. $D_{23} = \infty$) | $d_1 = 0$ |
| $D_{13} \geq D_{12} + D_{23}$ (incl. $D_{13} = \infty$) | $d_2 = 0$ |
| $D_{12} \geq D_{13} + D_{23}$ (incl. $D_{12} = \infty$) | $d_3 = 0$ |
| $D_{12} = D_{13} = \infty$ | $d_1 = \infty$ |
| $D_{12} = D_{23} = \infty$ | $d_2 = \infty$ |
| $D_{13} = D_{23} = \infty$ | $d_3 = \infty$ |
| $D_{12} = D_{13} = D_{23} = \infty$ | At least two of $d_1, d_2, d_3$ are infinite |

By comparison, for the CFN model considered in this paper, the analogous distance estimate (see, e.g., [37]) is

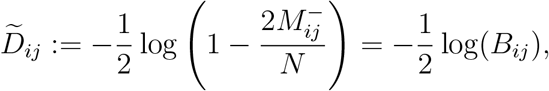

with the convention that 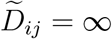 if *B*_*ij*_ *≤*0.

An inspection of Tables 1 and 2 reveals that, modulo the aforementioned change to the distance estimates to account for the different substitution models, the results of Theorem 2.4 coincide precisely with the predictions made in [28] for all data satisfying assumptions **A.1** and **A.2**. Thus, Theorem 2.4 puts the predictions in [28] on a sound theoretical basis, as it proves that certain properties of the maximum likelihood estimate are fully determined by whether or not the *data* satisfies the semialgebraic constraints of the model (i.e., the inequalities of Eqs. (1.16) and (1.17), which are mirrored by the inequality conditions in Tables 1 and 2).

In addition, Theorem 2.4 provides a characterization and better understanding of the way that the maximum likelihood estimator can fail to return a tree with biologically plausible branch lengths, but instead returns an estimate with edge parameters *θ* ∉ (0, 1)^3^.

Consider the data point

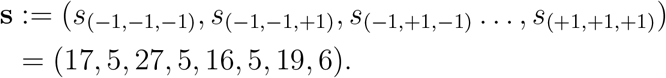

In [23], it was shown using algebraic methods that maximum likelihood fails to return an estimate with biologically plausible edge parameters for this data point. Instead, it was shown that for this data, the likelihood is maximized as the branch length of leaf 2 goes to infinity. It is easy to verify this conclusion with Theorem 2.4. Observe that

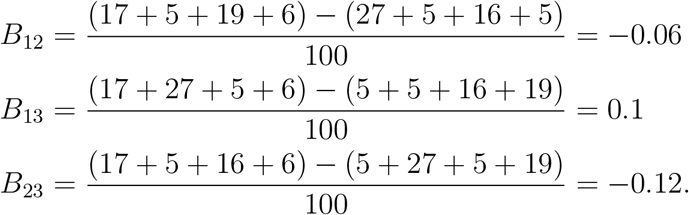

Since *B*_12_, *B*_23_ *<* 0, it follows by row 6 of Table 1 that the likelihood is maximized when

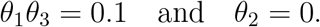

This agrees with the result in [23], as *θ*_2_ = 0 if and only if *d*_2_ = + ∞. Indeed, Theorem 2.4 characterizes all the ways that the likelihood estimate might be maximized when one or more branch lengths tend to infinity, up to our simplifying assumptions **A.1** and **A.2**.

Further, Theorem 2.4 provides a succinct explanation of why, for this data, it would be unreasonable to expect maximum likelihood to return a tree with biologically plausible edge parameters. For any tree with edge parameters *θ* ∈ (0, 1)^3^, the Fourier coordinates must satisfy the semi-algebraic constraint in Eq. (1.17), but for this data point, since *B*_12_, *B*_23_ *<* 0, the estimates of the Fourier coordinates fail to satisfy a corresponding positivity inequality.

The previous example elucidates one of the ways that long branches on a species tree can result in the MLE returning a boundary case: when data comes from a species tree with one or more very long branches relative to the size of *N*, it is more likely that one or more components of **B** will be negative, so that **B** ∉ 𝒟and hence the MLE must be on the boundary.

Nonetheless, this is not the full story. The next example shows how the MLE can lie on the boundary of the model even if all of the distance estimates are finite and positive. Consider the data

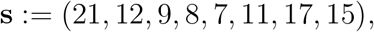

so that

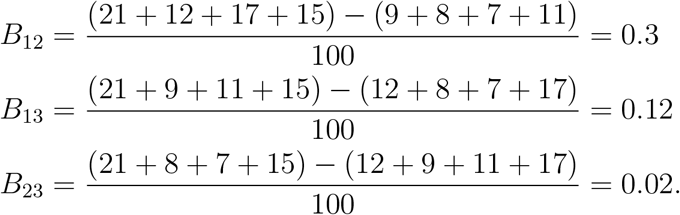

In this case *B*_*ij*_ *>* 0 for all *i, j* ∈[3], so the analogous inequality to Eq. (1.16) is satisfied: based on the data, all of the pairs of nucleotides are positively correlated, so no infinitely-long branches are to be expected; the distance estimates 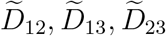 are all finite and positive.

However, by Theorem 2.4, this is not sufficient to guarantee that maximum likelihood will return a tree with biologically plausible parameters, since there is another semi-algebraic constraint (i.e., Eq. (1.17)) which must be satisfied for such trees. Indeed, since

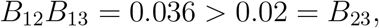

it follows that **B** ∉ 𝒟, and hence the data falls into case (i) of Theorem 2.4. More specifically, since this data corresponds to row 2 of Table 1, it follows that the likelihood is maximized when

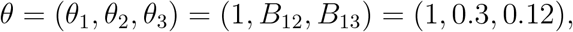

which does not correspond to a binary tree because the branch length of leaf 1 is zero.

One final and more general takeaway from Theorem 2.4 is that it highlights how the geometry of the statistical model, here determined by the semi-algebraic constraints Eqs. (1.16) and (1.17), influences the possible behavior of maximum likelihood estimation. Maximum likelihood returns a tree with biologically plausible branch lengths if and only if the data satisfies analogues of the polynomial inequalities Eqs. (1.16) and (1.17). In addition, when the data does *not* lie in the interior of the model, the question of which inequalities are not satisfied determines the different ways that maximum likelihood fails (i.e., which branches have lengths zero or infinity). This suggests that for phylogenetic trees with more than three leaves, a better understanding of the role that semialgebraic constraints play in maximum likelihood estimation may turn out to be useful in explaining some of the more complex behaviors of maximum likelihood estimation of phylogenetic trees. Of special interest to note are 4-leaf trees, due to the possibility of long-branch attraction. Indeed, recent work has shown that in the 4-leaf case, the study of semi-algebraic constraints for phylogentic models involves surprising subtleties that may be important for inference [6].

To be sure, in the cases of trees with *n* ≥ 4 leaves, to say nothing of more realistic and complicated substitution models, the increased algebraic complexity of the likelihood equations presents formidable obstacles. First, as shown in [20], when *n* ≥ 4, the likelihood equations do not have solutions which can be expressed in terms of pairwise sequence comparisons (as was done here and in [20]). Moreover, in many cases, closed form solutions are unlikely to exist at all; an example of this can be found in the analysis of the 4-leaf MC-comb in [10], where it was shown that the critical points of the likelihood function correspond to zeros of a degree 9 polynomial which cannot be solved by radicals.

Despite these limitations, solutions can nonetheless be obtained using tools from numerical algebraic geometry which return theoretically correct solutions with probability one (see, e.g., [23, 16, 9, 10]). Moreover, the *number* of solutions, called the *maximum likelihood (ML) degree*, can be computed using Gröbner basis techniques [21] and methods from singularity theory [30, 26].

## 4. Proof of the main result

Our proof of Theorem 2.4 considers the problem of maximizing the log-likelihood separately two cases:

1. the *interior case*, i.e., the problem of maximizing *ℓ* over all *θ* ∈ (0, 1)^3^, and
2. the *boundary cases*, corresponding to when *θ* ∈*∂*(0, 1)^3^.

For the boundary cases, we follow the approach taken by the authors of [28], who analyzed the maximum likelihood problem in the context of the Jukes-Cantor model and obtained analytic solutions for boundary cases there by decomposing the boundary ∂(0, 1)^3^ into 26 components, consisting of 8 vertices, 12 edges, and 6 faces, and then maximizing *ℓ* on each of those individually. Our approach is similar, though we group certain edges and faces together in those cases in which the analysis is similar.

The approach taken to proving Theorem 2.4 is as follows. First, the problems of maximizing *ℓ* in the interior case and boundary cases are considered separately in Sections 4.1 and 4.2 respectively. Section 4.3 presents several lemmas which compute and compare log-likelihoods of local maxima in various cases, with results summarized in Table 3. Finally, in Section 4.4, we utilize these results to prove Theorem 2.4.

**Table 3.** Summary of the results of Lemma 4.2, 4.3, 4.4, 4.6, and 4.5.

| Set $S$ | Criteria for $S \neq \emptyset$ | $\ell(\theta), \theta \in S$ |
| --- | --- | --- |
| $\mathcal{E}_{\text{int}}$ | $(B_{12}, B_{13}, B_{23}) \in \mathcal{D}$ | Eq. (4.3) |
| $\mathcal{E}_{\text{triv}}$ | always nonempty | $-\infty$ |
| $\mathcal{E}_{\text{ind}}$ | always nonempty | $-N \log 8$ |
| $\mathcal{G}_{\pi} (\pi \in \text{Alt}(3))$ | $B_{\pi} \in (0,1)$ | Eq. (4.23) |
| $\mathcal{F}_{\pi} (\pi \in \text{Alt}(3))$ | $B_{\pi'}, B_{\pi''} \in (0,1)$, where $\{\pi', \pi''\} = \text{Alt}(3) \setminus \{\pi\}$ | Eq. (4.18) |

### 4.1. Maximizing the log-likelihood on (0, 1)^3^

In this subsection we consider the problem of maximizing Eq. (1.8) on the set (0, 1)^3^. This set corresponds to those trees whose branches are of finite and nonzero length, when measured in expected number of mutations per site. Since (0, 1)^3^ is open, the existence of a local maximum is not guaranteed. The main result of this subsection gives, for generic positive data, necessary and sufficient conditions for *ℓ* to have a local maximum in (0, 1)^3^, and a formula if it exists; it also shows *ℓ* has at most one local maximum on (0, 1)^3^.

We begin with an important definition and a technical lemma.

Let 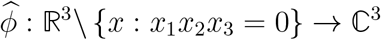 be defined by

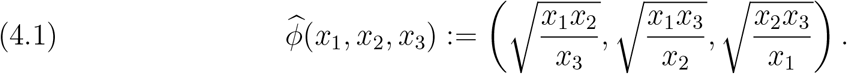

Further, let *ϕ* be the restriction of 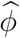 to 𝒟:

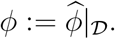

The next lemma summarizes some useful properties of *ϕ*.

#### Lemma 4.1.

The function *ϕ* : 𝒟 *→* (0, 1)^3^ is a continous bijection, with inverse function *ϕ*^*−*1^ : (0, 1)^3^ *→*𝒟 given by

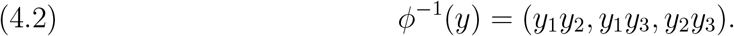

*Proof of Lemma 4*.*1*. First, note that if *x* = (*x*_1_, *x*_2_, *x*_3_) ∈𝒟 then *ϕ*(*x*) ∈(0, 1)^3^ by the definition of 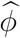 and 𝒟. Moreover, since *x*_1_, *x*_2_, *x*_3_ ≠ 0, whenever *x* = (*x*_1_, *x*_2_, *x*_3_) ∈𝒟, it follows that *ϕ* is continuous on 𝒟.

Next, to show that *ϕ* is injective, let 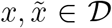 such that 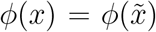. Let *ϕ*_1_ and *ϕ*_2_ denote the first and second components of *ϕ* respectively. Then 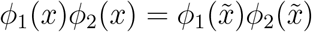 or equivalently, 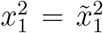. Since 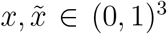, this implies 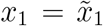. Similar arguments show that 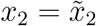 and 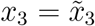, and hence 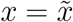. Therefore *ϕ* is injective.

Next we show that *ϕ* is surjective. Let *y* ∈ (0, 1)^3^ be arbitrary. Then it is easy to see that the point 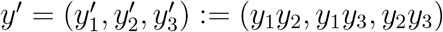 satisfies

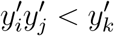

for all choices of distinct *i, j, k* ∈ [3], and hence *y*^*′*^∈𝒟. Moreover, *ϕ*(*y*^*′*^) = *y* by definition of 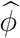. Therefore *ϕ* is surjective and has an inverse given by the formula *ϕ*^*−*1^(*y*) = *y*^*′*^, which is precisely the formula in Eq. (4.2). □

We are now ready to state the main lemma of this subsection, which solves the problem of maximizing *ℓ* on the set (0, 1)^3^, or in other words, solves the maximum likelihood problem for biologically plausible parameters.

#### Lemma 4.2

(MLE for 3 Leaf Tree – Interior Case). Assume that **A.1** and **A.2** hold, and let

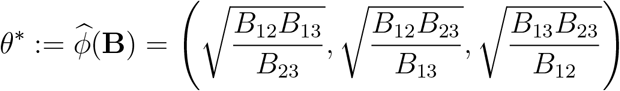

If **B** ∈𝒟 then *θ*^*\**^is the unique local maximum of *ℓ* in (0, 1)^3^ and has log-likelihood

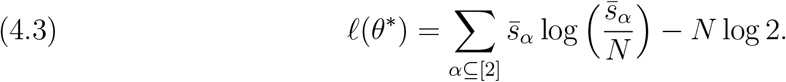

On the other hand, if **B ∉** *D* then *ℓ* has no local maximum on (0, 1)^3^.

*Proof of Lemma 4*.*2*. For ease of notation, we will write

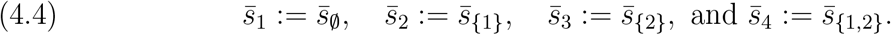

It follows by Eq. (1.8) and Eq. (1.11) that

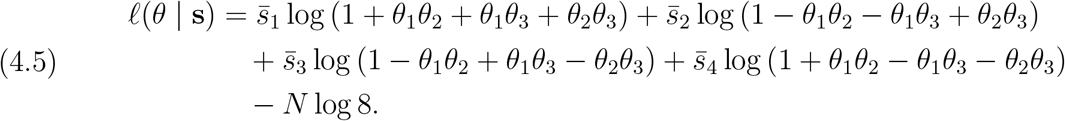

An initial attempt to compute the critical points of *ℓ*(*θ*) directly by taking the gradient of *ℓ* yields a polynomial system which is difficult to solve analytically, so instead we modify this approach by first considering a different function whose extrema are closely related to those of *ℓ*.

To define this function, first let 𝒟_*F*_ *⊆ ℝ*^3^ be the intersection of half-spaces defined by the following inequalities

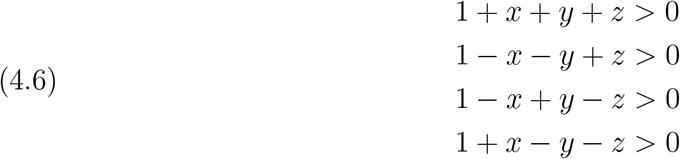

and let *F* : 𝒟_*F*_ *→ℝ* be defined by

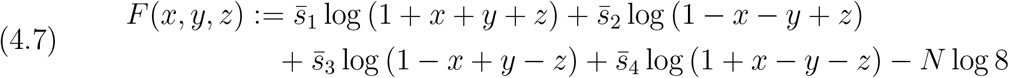

The significance of *F* is owed to the observation that *ℓ* = *F °ϕ*^*−*1^, which is proved in the next claim.

#### Claim 1

For all *θ* ∈ (0, 1)^3^,

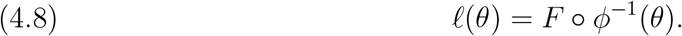

*Proof of Claim 1*. Since the domain of *ℓ* is (0, 1)^3^, it follows from Lemma 4.1 and Eqs. (4.5) and (4.7) that Eq. (4.8) holds provided that *F* is defined on the image of *ϕ*^*−*1^. Therefore, since im(*ϕ*^*−*1^) = 𝒟 and dom(*F*) = 𝒟_*F*_ it will suffice to show that 𝒟 *⊆𝒟*_*F*_.

Let *u ∈𝒟*. Then by Lemma 4.1 there exist *w*_1_, *w*_2_, *w*_3_ ∈ (0, 1) such that

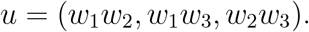

To show that *u ∈ D*_*F*_, it suffices by Eq. (4.6) to show that

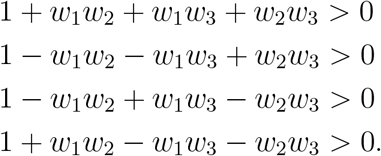

The first of these four equations holds trivially. As for the other three, we will only prove 1 + *w*_1_*w*_2_ *− w*_1_*w*_3_ *− w*_2_*w*_3_ *>* 0, as the other two inequalities can be proved in the same manner. Let *h*(*w*_1_, *w*_2_) := 1 + *w*_1_*w*_2_ *− w*_1_ *− w*_2_. Since *w*_1_, *w*_2_ *>* 0 and *w*_3_ *<* 1, it holds that

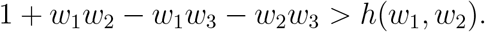

Therefore it will suffice to show that *h*(*w*_1_, *w*_2_) *≥* 0 for all *w*_1_, *w*_2_ ∈ [0, 1]. Indeed, using calculus it is easy to see that *h* is minimized on [0, 1] *×* [0, 1] when at least one of the arguments *w*_1_, *w*_2_ equals one, and that the minimum is zero. Therefore 1 + *w*_1_*w*_2_ *−w*_1_*w*_3_ *− w*_2_*w*_3_ *>* 0. We conclude that 𝒟 ⊆𝒟_*F*_, as required to prove the claim. □

The next two claims serve to characterize the extrema of *F*.

#### Claim 2.

The point **B** = (*B*_12_, *B*_13_, *B*_23_) is the unique critical point of *F*.

*Proof of Claim 2*. It is first necessary to verify that **B** is in the domain of *F*. To do this, it will suffice to show that **B** satisfies the inequalities in Eq. (4.6).

Using Eq. (2.2) and the observation 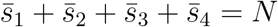, it is easy to check that

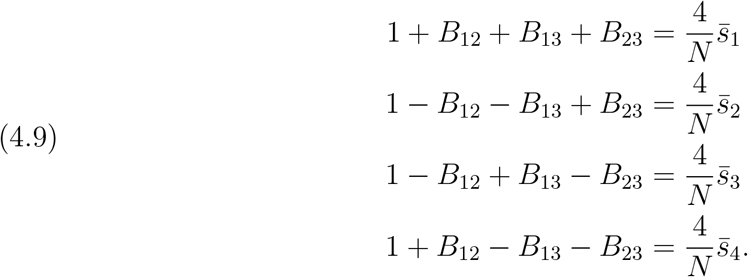

Therefore by **A.1**, it follows that **B** = (*B*_12_, *B*_13_, *B*_23_) satisfies the inequalities in Eq. (4.6), as required. Therefore **B** is in the domain of *F*.

We now proceed with a standard critical point calculation. Letting *v*_1_ = (1, 1, 1)^⊤^, *v*_2_ = (*−*1, *−*1, 1)^⊤^, *v*_3_ = (*−*1, 1, *−*1)^⊤^, *v*_4_ = (1, *−*1, *−*1)^⊤^, and taking partial derivatives of *F* in Eq. (4.7) with respect to the variables *x, y* and *z*, it follows that for all *u* = (*x, y, z*)^⊤^ ∈𝒟_*F*_,

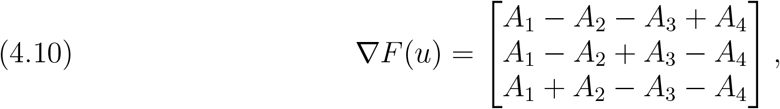

where

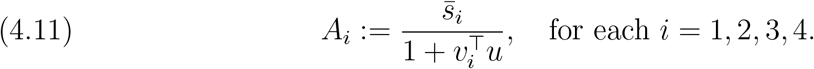

Setting *∇F* = 0 and using Eq. (4.10), we deduce that the following system of equations holds:

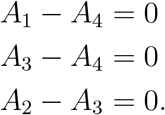

Substituting the formulas for the *A*_*i*_’s from Eq. (4.11) and rearranging terms, we obtain

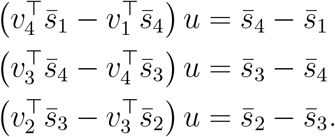

Writing this out, this is the matrix equation

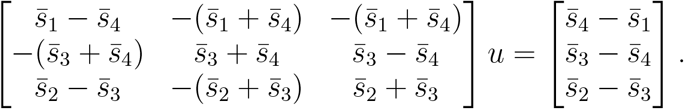

Using the fact that 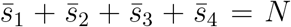, one can check that the 3 *×* 3 matrix in the above equation has inverse

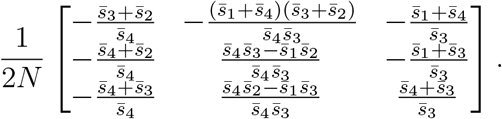

Therefore

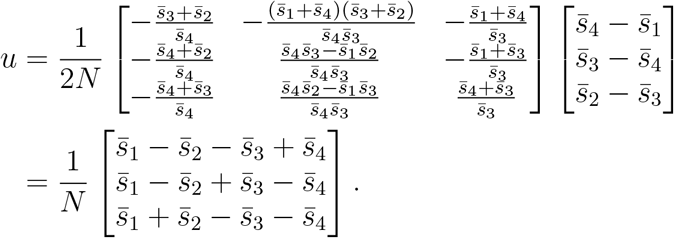

By Eq. (2.2), the right-hand side is precisely the vector (*B*_12_, *B*_13_, *B*_23_)^⊤^, and therefore we conclude that **B** = (*B*_12_, *B*_13_, *B*_23_) is the unique critical point of *F* on its domain 𝒫. This completes the proof of the claim. □

Let *H*_*F*_ denote the Hessian matrix of *F* ; that is,

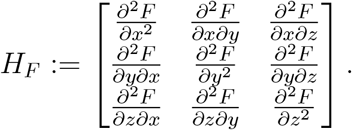

It is clear that *F* is twice-differentiable on its domain, so *H*_*F*_ (*x*) is defined for all *x* ∈𝒟_*F*_.

#### Claim 3.

*H*_*F*_ (**B**) is negative definite.

*Proof of Claim 3*. Since *H*_*F*_ (**B**) is a real symmetric matrix, it will suffice to show that its eigenvalues are all negative, as this will imply that *H*_*F*_ (**B**) is negative definite.

Using the code in Appendix B, we first compute the characteristic polynomial of *H*_*F*_ :

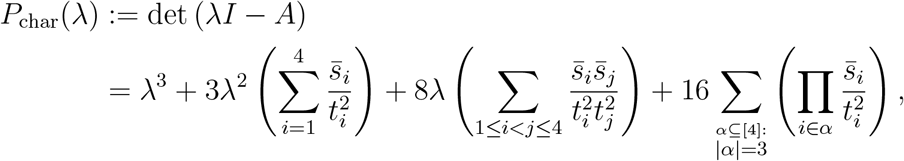

where *t*_1_ = 1 + *x* + *y* + *z, t*_2_ = 1 − *x − y* + *z, t*_3_ = 1 − *x* + *y − z, t*_4_ = 1 + *x − y − z*, and 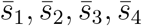 are defined in Eq. (4.4).

We need to show that *P*_char_ has only negative roots, as this will imply that all three eigenvalues of *H*_*F*_ are negative, and hence that *H*_*F*_ (**B**) is negative definite. Since *H*_*F*_ is a real symmetric matrix, the roots of *P*_char_ are all real numbers, and therefore it will be enough to show that *P*_char_ has no nonnegative roots. Indeed, since all the coefficients of *P*_char_ are positive, Decartes’ rule of signs implies that *P*_char_ has no positive roots. Moreover since 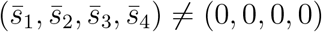 by **A.1**, the constant term in *P*_char_ is nonzero, and hence *P*_char_(0) = 0 as well. We conclude that *P*_char_ has no nonnegative roots, as required to prove the claim. □

Using the results of the previous three claims, the next two claims will together characterize the local maxima of *ℓ* on (0, 1)^3^.

#### Claim 4

If **B** *∈𝒟* then *θ*^*\**^ ∈ (0, 1)^3^ and *θ*^*\**^ is a local maximum of *ℓ*.

*Proof of Claim 4*. Observe that *F* and *ϕ*^*−*1^ are both differentiable on their respective domains. Therefore if *θ* ∈ (0, 1)^3^ and *x* = *ϕ*^*−*1^(*θ*), then using the chain rule to differentiate Eq. (4.8) gives

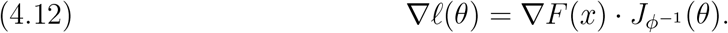

Suppose **B** ∈**𝒟**. Then by Lemma 4.1, *θ*^*\**^ = *ϕ*(**B**) ∈ (0, 1)^3^ and **B** = *ϕ*^*−*1^(*θ*^*\**^). Therefore Eq. (4.12) implies

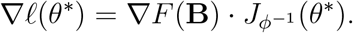

By Claim 2, **B** is a critical point of *F*, i.e., *∇F* (**B**) = 0. Therefore

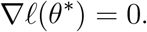

This shows that *θ*^*\**^ is a critical point of *ℓ*.

Next we show that *θ*^*\**^ is a local maximum of *ℓ*. Since *ℓ* = *F* ○ *ϕ*^*−*1^ by Eq. (4.8), and since *F* and *ϕ*^*−*1^ are both twice differentiable on their respective domains, therefore it follows by the chain rule for Hessian matrices (see, e.g., [24, p.125-126]), that *ℓ* is also twice differentiable and has Hessian matrix

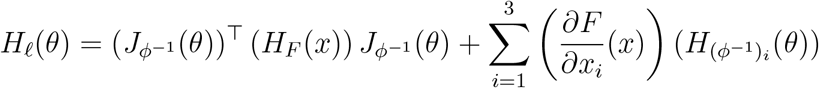

where

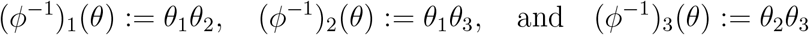

for all *θ* ∈ (0, 1). Since **B** is a critical point of *F* due to Claim 2, we have 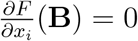 for each *i* = 1, 2, 3. Therefore

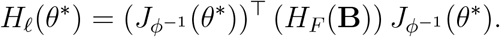

Since *H*_*F*_ (**B**) is a negative definite matrix by Claim 3, and since det 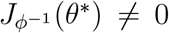 by Eq. (4.13), we conclude that *H*_*ℓ*_(*θ*^*\**^) and *H*_*F*_ (**B**) are similar matrices, and hence *H*_*ℓ*_(*θ*^*\**^) is negative definite as well. Therefore by the second derivative test (see, e.g., [24, Theorem 4, p.140]), the point *θ*^*\**^ is a local maximum of *ℓ*. This completes the proof of the claim. □

#### Claim 5

If *θ* ∈ (0, 1)^3^ is a local maximum of *ℓ* then **B** *∈𝒟* and *θ* = *θ*^*\**^.

*Proof of Claim 5*. Let *θ* ∈ (0, 1)^3^ be a local maximum of *ℓ* and let *x* = *ϕ*^*−*1^(*θ*). Since *θ* a critical point, Eq. (4.12) implies

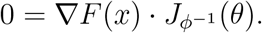

Since

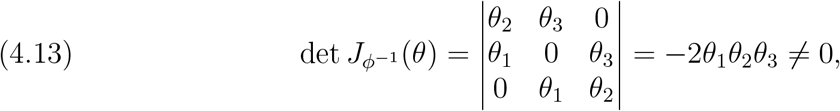

it follows that 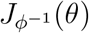 is non-singular, and hence

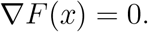

Therefore *x* is a critical point of *F*. Since the only critical point of *F* is **B** by Claim 3, it follows that *x* = **B**. Since *x ∈𝒟* by Lemma 4.1, this implies that **B** *∈𝒟*. Therefore

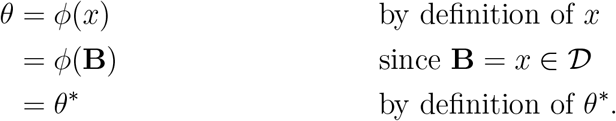

This completes the proof of the claim. □

We can now use Claims 4 and 5 to prove the theorem. If **B** ∈𝒟then Claims 4 and 5 imply that *θ*^*\**^ is the unique local maximum in (0, 1)^3^. On other other hand, if **B ∉ 𝒟**, then the contraposition of Claim 5 implies *ℓ* has no local maximum in (0, 1)^3^. This proves the first part of Lemma 4.2; it remains only to prove Eq. (4.3).

Indeed, if **B** *∈𝒟* then since *θ*^*\**^= *ϕ*(*B*_12_, *B*_13_, *B*_23_), it follows by definition of *ϕ* that

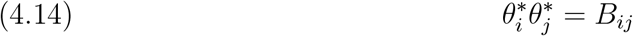

for all *i, j* ∈ [3] such that *i < j*. Plugging Eq. (4.14) into Eq. (4.9),

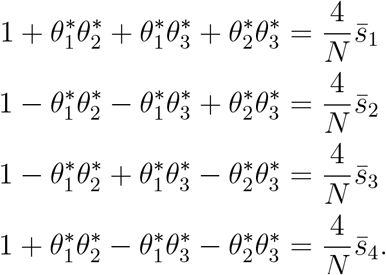

Therefore by Eq. (4.5),

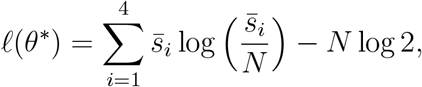

which is precisely Eq. (4.3). This completes the proof of Lemma 4.2. □

### 4.2. Maximizing the log-likelihood on ∂(0, 1)^3^

In this subsection we consider the problem of maximizing *ℓ* on the boundary ∂(0, 1)^3^. As discussed at the beginning of Section 4, this corresponds to the boundary of the unit cube, consisting of 6 faces, 12 edges, and 8 vertices. The lemmas in this section consider the problem of maximizing *ℓ* on various groupings of these components.

The eight vertices of the unit cube are simply the elements of the set {0, 1} ^3^. The twelve edges are the sets

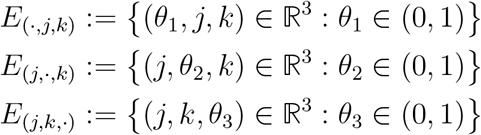

defined for *j, k* ∈{0, 1}. The 6 faces are the sets of the form

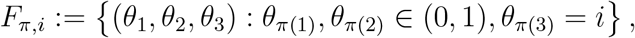

where *π* ∈Alt(3) and *i* ∈{0, 1}.

The task at hand is to maximize *ℓ* on each of these boundary sets. We begin with the next lemma, which utilizes assumption **A.1** to dispatch the edge and vertex boundary cases which are “trivial” in the sense of never containing the maximum.

#### Lemma 4.3

(Trivial cases: ℰ_triv_). Assume that **A.1** holds. Let

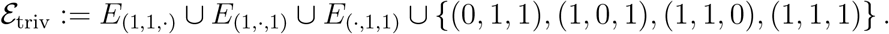

If *θ ∈ℰ*_triv_ then *ℓ*(*θ* | *s*) = *−∞*.

*Proof*. If *θ ∈ E*_(1,1,*·*)_ *∪E*_(1,*·*,1)_ *∪E*_(*·*,1,1)_ *∪*{(0, 1, 1), (1, 0, 1), (1, 1, 0), (1, 1, 1)*}* then there exist two leaves *i, j*∈[3] with *i ≠ j* such that *θ*_*i*_ = *θ*_*j*_ = 1. By **A.1**, there exists a *σ* ∈{ 1, *−*1} ^3^ with *σ*_*i*_ *≠ σ*_*j*_ and *s*_*σ*_ *>* 0. Moreover, by Eq. (1.2), the probability of a transition occurring along the path between leaves *i* and *j* is zero. In particular, since ℙ [*X* = *σ*] ≤ ℙ [*X*_*i*_ *≠ X*_*j*_], this implies that ℙ [*X* = *σ*] = 0. Therefore

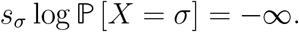

Therefore *ℓ*(*θ*) = *−∞* by Eq. (1.8). □

The next lemma considers the remaining 4 vertices of the unit cube (1, 0, 0), (0, 1, 0), (0, 0, 1), and (0, 0, 0), as well as the edges *E*_(0,0,*·*)_, *E*_(0,*·*,0)_, and *E*_(*·*,0,0)_. The union of these boundaries is the following set:

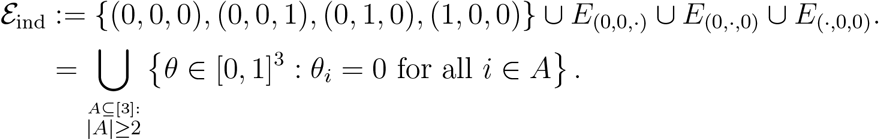

The next lemma shows that ℰ_ind_ consists of all choices of numerical parameter values under which the *X*_1_, *X*_2_, and *X*_3_ are independent.

#### Lemma 4.4

(Log-likelihood of *ℓ* on ℰ_ind_). If *θ ∈ℰ*_ind_ then

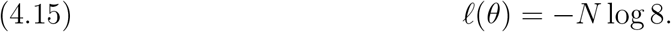

*Proof*. Suppose *θ* ∈ℰ_ind_, and let *i, j*∈ [3] such that *i ≠ j*. Then the path between leaves *i* and *j* contains an edge *e* such that *θ*_*e*_ = 0. Therefore by Lemma 1.3, the random variables *X*_1_, *X*_2_ and *X*_3_, are mutually independent. Therefore for all *σ ∈ {−*1, 1*}*^3^,

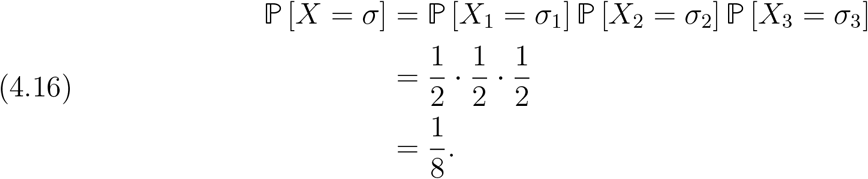

Substituting this into Eq. (1.8) implies Eq. (4.15). □

The next lemma of this section considers the problem of maximzing *ℓ* on the three faces *F*_*π*,1_, *π* ∈Alt(3). An example of the graphical model correponding to these cases is shown in Fig. 3.

#### Lemma 4.5

(Maximizers of *ℓ* on *F*_*π*,1_). Let *π* ∈Alt(3), and let *θ* ∈*F*_*π*,1_. Then the set of local maxima of *ℓ* on *F*_*π*,1_ is

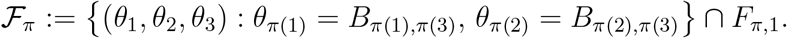

Moreover, if *θ ∈ F*_*π*_ then

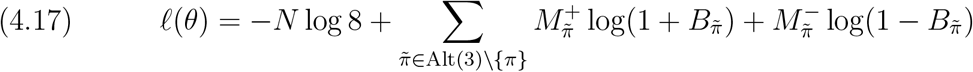

or equivalently (4.18)

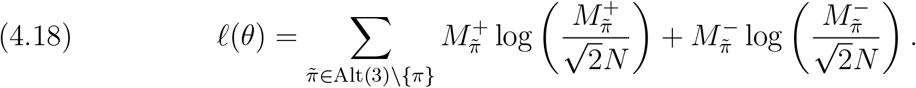

*Proof*. Let *π* ∈Alt(3), and let *i* = *π*(1), *j* = *π*(2), and *k* = *π*(3). Let *θ*_*i*_, *θ*_*j*_ ∈(0, 1) be arbitrary. Since *θ*_*k*_ = 1, it follows by Eq. (1.12)

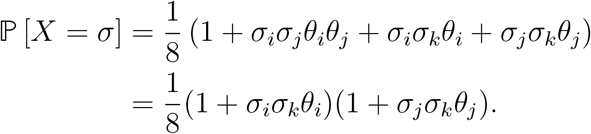

Hence Eq. (1.8) can be written as

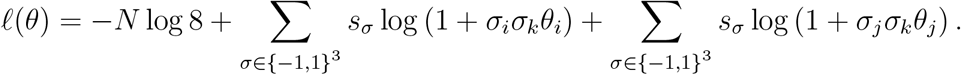

Therefore

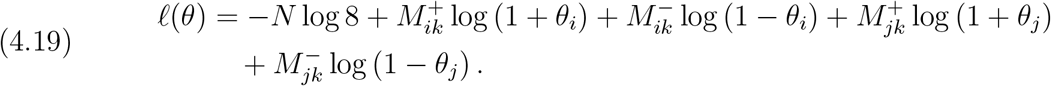

Differentiating with respect to *θ*_*i*_ and *θ*_*j*_, it follows that for each *u ∈{i, j}*,

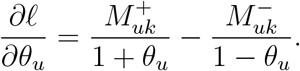

Solving the system *∇ℓ*(*θ*) = 0, we obtain the solution satisfying

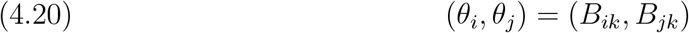

and if *B*_*ik*_, *B*_*jk*_ ∈ (0, 1) then Eq. (4.20) determines the unique critical point of *ℓ* on the set *F*_*π*,1_; on the other hand, if *B*_*ik*_ ∉ (0, 1) or *B*_*jk*_ *∉* (0, 1), then *ℓ* has no critical points on *F*_*π*,1_.

Moreover, this critical point on *F*_*π*,1_ is a maximum by the second derivative test since for all *θ ∈ F*_*π*,1_, the Hessian matrix

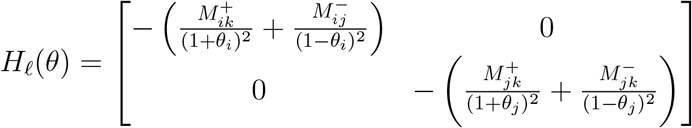

is negative definite.

Finally, if Eq. (4.20) holds, plugging *θ*_*i*_ = *B*_*ik*_ and *θ*_*j*_ = *B*_*jk*_ into Eq. (4.19) gives

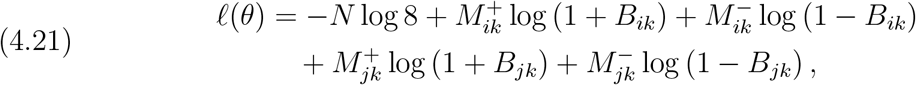

which is nothing but Eq. (4.17) written in a different notation. It remains to prove Eq. (4.18). Observe that 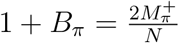 and 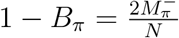 for all *π* ∈ Alt(3). Eq. (4.18) can be obtained by making these substitutions in Eq. (4.17) and then simplifying using logarithm properties and 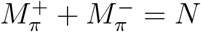. □

The final lemma of this section considers the problem of maximizing *ℓ* when exactly one parameter in *{θ*_1_, *θ*_2_, *θ*_3_*}* is assumed to be zero. This pertains to each of the 3 remaining faces *F*_*π*,0_, *π* ∈ Alt(3), as well as the remaining six edges *E*_(*·*,0,1)_, *E*_(*·*,1,0)_, *E*_(0,*·*,1)_, *E*_(1,*·*,0)_, *E*_(0,1,*·*)_, and *E*_(1,0,*·*)_. Such boundaries represent one of two graphical models, like those shown in Fig. 2. We group the boundary cases up into the following three sets, defined for each *π* ∈ Alt(3):

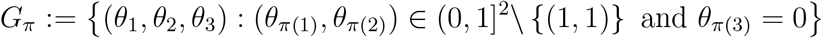

**Figure 2.**
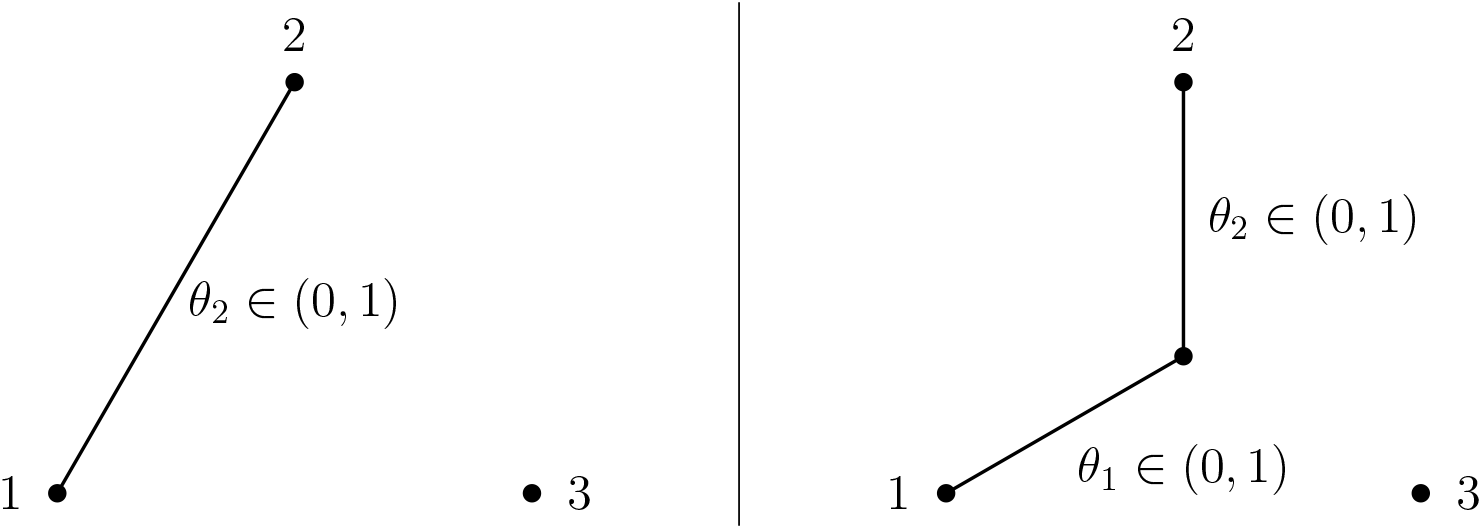
The graphical model corresponding to a 3-leaf “tree” with edge parameters *θ* ∈*E*_(1,*·*,0)_ (left) or *θ* ∈*F*_(*·,·*,0)_ (right). In both cases, the biological meaning of *θ*_3_ = 0 is that species 3 is “infinitely far away” from species 1 and 2, when distances are measured in expected number of mutations per site. By Lemma 1.3, (*X*_1_, *X*_2_) ╨ *X*_3_, and for this reason we depict vertex 3 as a disconnected vertex.

**Figure 3.**
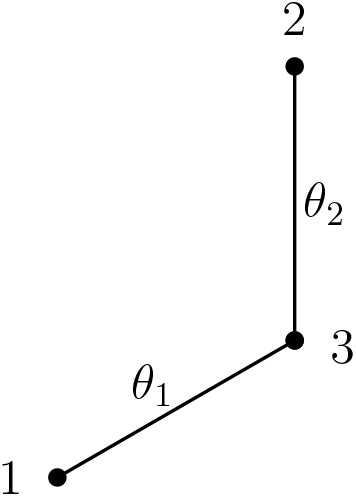
An example of a tree with numerical parameters *θ* ∈*F*_*π*,1_ = *θ* : *θ*_1_, *θ*_2_ (0, 1), *θ*_3_ = 1 with *π* = (1) the identity permutation. Since *θ*_3_ = 1, Eq. (1.2) implies that no transitions can occur on leaf 3, and hence vertex 3 is regarded to lie on the path between leaves 1 and 2.

Interpreted geometrically, *G*_*π*_ consists of the union of one face and two of its adjacent edges:

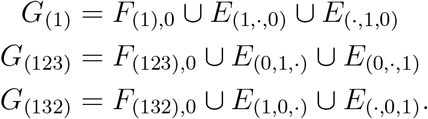

#### Lemma 4.6

(Maximizers of *ℓ* on *G*_*π*_). Let *π* ∈Alt(3). The local maxima of *ℓ* on *G*_*π*_ are the points in the set

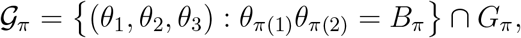

all of which have log-likelihood

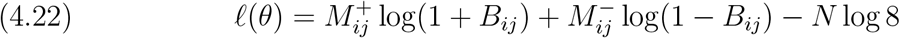

or equivalently

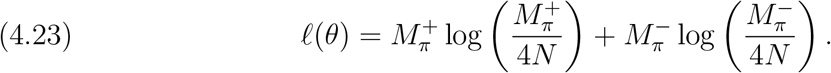

In particular, this implies that if *B*_*π*_ *∉* (0, 1) then *ℓ* has no local maxima on *F*.

The proof of this lemma is similar to that of Lemma 4.5, and can be found in Appendix A.

The key results of Sections 4.1 and 4.2 are summarized in Table 3. In that table, computed are the maximizer(s) of *ℓ* on the interior and boundary of the unit cube (namely, the sets *E*_int_, *ℰ*_triv_, *ℰ*_ind_, *G*_*π*_, and *F*_*π*,1_, *π* ∈ Alt(3), which together partition the closed unit cube). The corresponding sets of maximizers on each of these sets (ℰ_int_, *ℰ*_triv_, *ℰ*_ind_, *𝒢*_*π*_, and ℱ_*π*_, *π* ∈ Alt(3), respectively) are level sets of *ℓ*, although some of them may be empty depending on the data. The necessary and sufficient conditions for each of them to be nonempty, as well as the value that the log-likelihood function takes on each of them, are shown in the second and third columns of the table.

### 4.3. Comparisons of likelihoods

To prove Theorem 2.4 will require some comparisons of the log-likelihoods of elements of ℰ_int_, *ℰ*_ind_, 𝒢_*π*_, and ℱ_*π*_, (*π* ∈ Alt(3)). In this subsection, we prove several lemmas toward this end.

We will make use of the information inequality, which we state next (for proof, see, e.g., Theorem 2.6.3 in [13]).

#### Theorem 4.7

(Information Inequality). Let *k ≥* 1. If 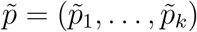 and 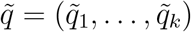 satisfy 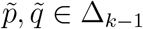then

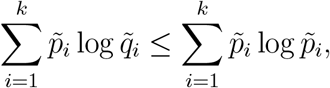

with equality if and only if 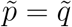.

The next two lemmas utilize the information inequality to show that elements of ℰ_int_ have greater log-likelihood than elements of ℱ_*π*_ and 𝒢_*π*_, for all *π* ∈ Alt(3).

#### Lemma 4.8

(ℰ_int_ vs ℱ_*π*_). If *θ*^*\**^ *∈ℰ*_int_ then

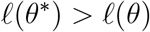

for all *θ ∈ℱ*_*π*_ and all *π* ∈ Alt(3).

*Proof*. We prove the case with *π* = (1) as the proofs for the cases with *π* ∈*{*(123), (132)} are similar.

Let *θ ∈ℱ*_*π*_. By Eq. (4.18),

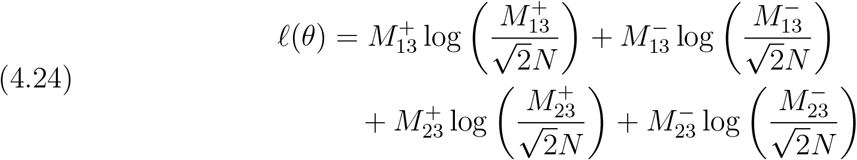

Observe that

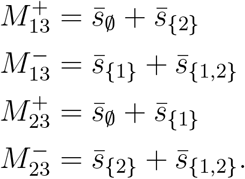

Therefore we can rewrite Eq. (4.24) as

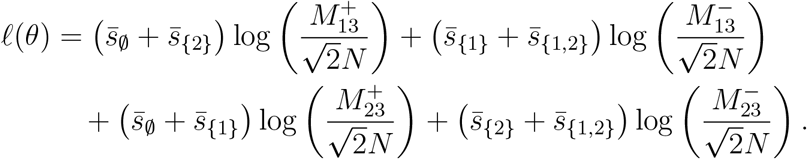

Regrouping terms gives

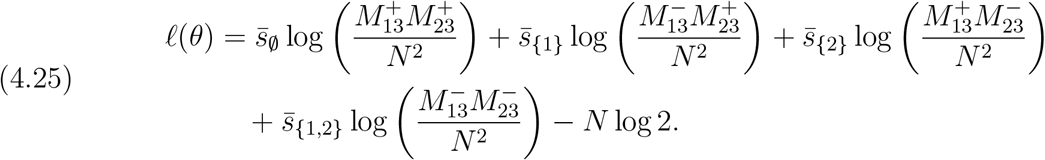

To apply Theorem 4.7, it is first necessary to verify that

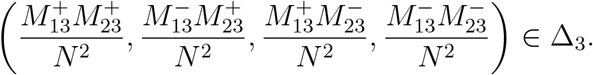

Since the entries of this vector are clearly nonnegative, it suffices to show that they sum to 1. Indeed, using 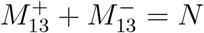 and 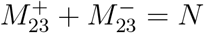, we have

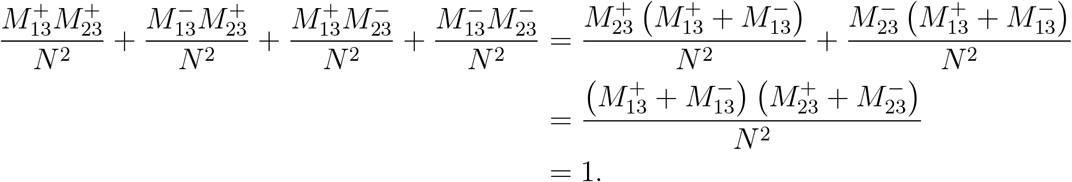

Therefore by applying Theorem 4.7 to the right hand side of Eq. (4.25),

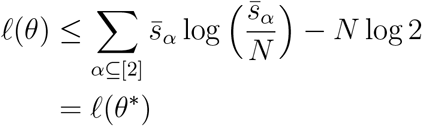

where the last equality follow from Eq. (4.3). □

A similar application of Theorem 4.7 can be used to prove the following theorem; the details of the proof can be found in Appendix A.

#### Lemma 4.9

(ℰ_int_ vs 𝒢_*π*_). Assume that **A.1** holds. If *θ*^*\**^ *∈ℰ*_int_ then

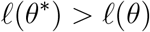

for all *θ ∈𝒢*_*π*_ and all *π* ∈ Alt(3).

The next lemma compares the likelihoods of elements in ℱ_*π*_ and 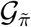 when 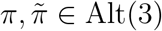 are distinct.

#### Lemma 4.10

(*ℱ*_*π*_ vs 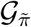 for 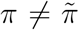). Assume that **A.1** holds. Let *π*_1_, *π*_2_ and *π*_3_ denote the three distinct elements of Alt(3). If 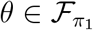 then

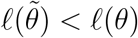

for all 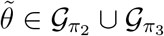.

*Proof*. Suppose 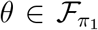, and suppose that 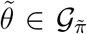 for some 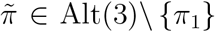. Without loss of generality, assume 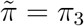. Then by Eqs. (4.17) and (4.22)

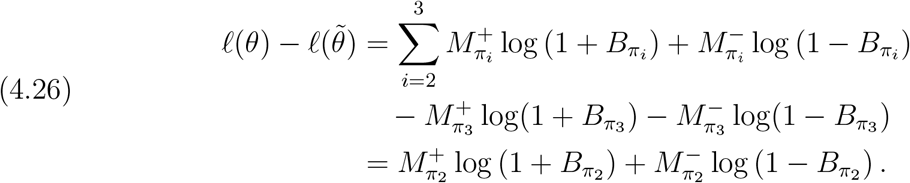

Since 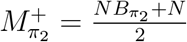 and 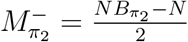, it follows that

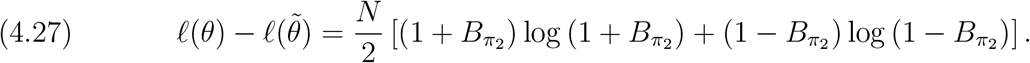

As a function of 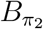, the right-hand side is strictly increasing on (0, 1), which can be seen by differentiating and observing that the derivative is positive on this interval. Moreover, we note that 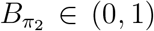, a fact which follows from the hypothesis that 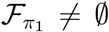 (see Table 3). Therefore, since the right-hand side of Eq. (4.27) is strictly increasing on (0, 1), and since 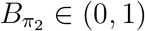, it follows that

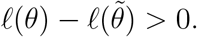

This completes the proof of the lemma. □

The next lemma compares the likelihoods of elements in ℱ_*π*_ and 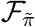 when 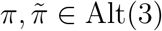 are distinct.

#### Lemma 4.11

(ℱ_*π*_ vs 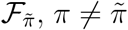). Let 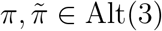 be distinct, and suppose that *θ ∈ℱ*_*π*_ and 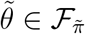. Then

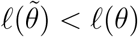

if and only if

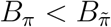

*Proof*. By Eq. (4.17)

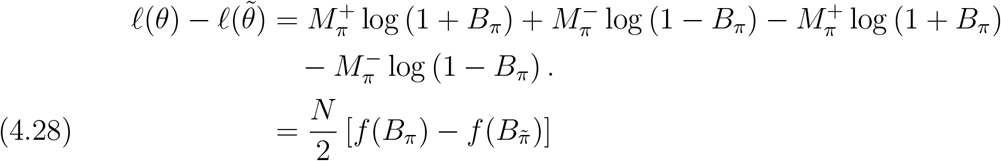

where *f* (*x*) := (1 + *x*) log (1 + *x*) + (1 *− x*) log (1 *− x*). Note that

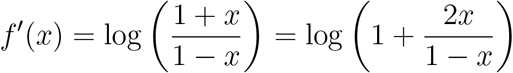

which is positive for all *x* ∈ (0, 1). Therefore *f* is increasing on (0, 1). Moreover, 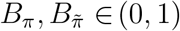 since 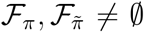 (see Table 3). Taken together, these facts along with Eq. (4.28) imply that 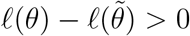 if and only if 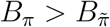. □

The next lemma compares the log-likelihods of elements in 𝒢_*π*_ and 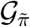, for distinct elements 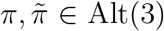. The proof is similar to that of Lemma 4.11 and can be found in Appendix A.

#### Lemma 4.12

(𝒢_*π*_ vs 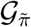 for 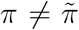). Let *π*, 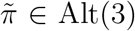 such that 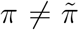, and let *θ ∈𝒢*_*π*_ and 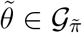. Assume that **A.1** holds. If

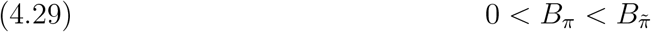

then

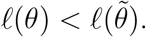

The next lemma shows that elements of 𝒢_*π*_, *π* ∈ Alt(3) have greater log-likelihood than elements in ℰ_ind_. The proof is straightforward and can be found in Appendix A.

#### Lemma 4.13

(𝒢_*π*_ vs ℰ_ind_). If *θ ∈𝒢*_*π*_ for some *π* ∈ Alt(3) then

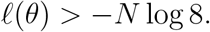

### 4.4. Final analysis

Using the lemmas from Sections 4.1, 4.2 and 4.4, we are now ready to prove Theorem 2.4.

*Proof of Theorem 2*.*4*. We claim that *θ ↦ ℓ*(*θ*) is upper semicontinuous on its domain [0, 1]^3^. To see this, first observe that the function *L* : [0, 1]^3^ *→* [0, *∞*) defined by

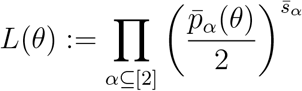

is continuous since for all 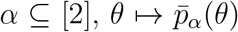 is a polynomial in the variables *θ*_1_, *θ*_2_, *θ*_3_ ∈ [0, 1] by Eq. (1.12). Therefore, since *ℓ* = log *L* and since log(·) is increasing and upper semicontinuous on [0, ∞), it follows that *ℓ* is upper semicontinuous on [0, 1]^3^. This proves the claim.

Since *ℓ* is upper semicontinuous, it has at least one maximizer on [0, 1]^3^. In order to find the maximizer(s), observe that since

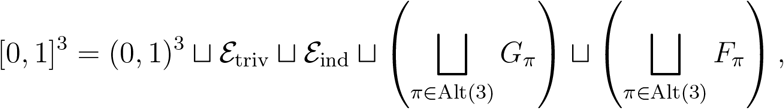

it suffices to consider the maximizers of *ℓ* on each of these sets. By Lemma 4.3,

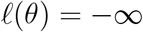

whenever *θ* ∈ *ℰ*_triv_. Therefore if 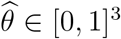 is a global maximum of 𝓁, then

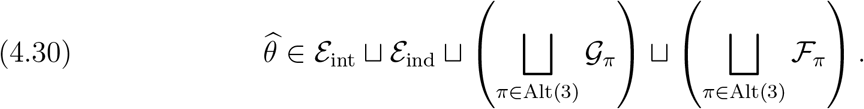

In Lemmas 4.2, 4.4, 4.6, and 4.5, we computed the log-likelihood of the points in each of the eight sets in this disjoint union (see Table 3 for a summary of these results), so the rest of the proof will simply be a comparison of the likelihoods of elements of these sets.

We start by proving that if **B** ∈ 𝒟, then the maximum is given by Eq. (2.3). Suppose **B** ∈ 𝒟. Then by Lemma 4.2, *ε*_int_ is nonempty and consists of a single element

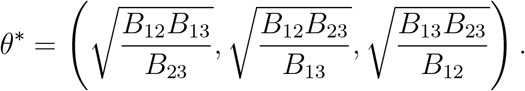

Moreover, Lemmas Lemma 4.8, 4.9, 4.10, and 4.13 together imply that *θ*^∗^ is the global maximizer of 𝓁. This proves Eq. (2.3).

Henceforth assume **B** ∉ 𝒟, so that *ℰ*_int_ = ∅ by Lemma 4.2. Therefore

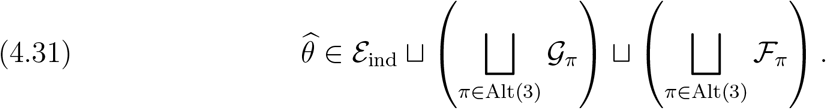

Next we will prove part (i) in the statement of the lemma. Suppose that 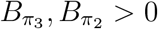. It will suffice to show that 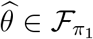.

By the criteria shown in Table 3, it holds that 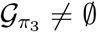 and 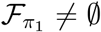. Let 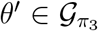 and 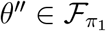. By Lemmas 4.12 and 4.13,

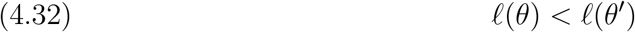

whenever 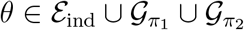. In addition, since *π*_1_ ≠ *π*_3_, Lemma 4.10 implies

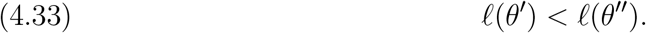

Therefore by Eqs. (4.31) to (4.33),

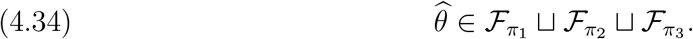

Finally, observe that by Lemma 4.11,

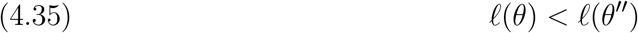

whenever 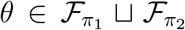. By Eqs. (4.34) and (4.35), we conclude that 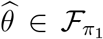. This proves part (i) of the lemma.

Next, suppose that 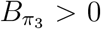 and 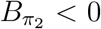. In order to prove part (ii) in the statement of the lemma, it will suffice to show that 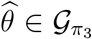. By the criteria in Table 3, ℱ _*π*_ = ∅ for all *π* ∈Alt(3), and 𝒢_*π*_ = ∅ for all *π* ∈ Alt(3)\{*π*_1_, *π*_2_*}*. Therefore by Eq. (4.31),

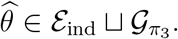

By Lemma 4.9, the elements of 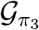 have strictly larger log-likelihood than the elements of ℰ_ind_. Therefore 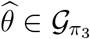. This proves part (ii) of the lemma.

It remains to prove part (iii) in the statement of the lemma. Suppose 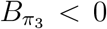. By the criteria in Table 3, ℱ_*π*_ = ∅ and 𝒢_*π*_ = ∅ for all *π* ∈ Alt(3). Therefore by Eq. (4.31), 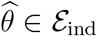. This proves part (iii), which completes the proof of the theorem.

## Acknowledgements

Rodriguez is partially supported by the Alfred P. Sloan Fellowship and the Office of the Vice Chancellor for Research and Graduate Education at UW-Madison with funding from the Wisconsin Alumni Research Foundation. Additional parts of this research were performed while JIR and MH were visiting the Institute for Mathematical and Statistical Innovation (IMSI) for the semester-long program on “Algebraic Statistics and Our Changing World,” IMSI is supported by the National Science Foundation (Grant No. DMS-1929348). MH is grateful for helpful discussions and feedback from Cécile Ané and Kaie Kubjas. The authors are thankful for the valuable comments by the referees.

## Appendix A. Omitted proofs

### A.0.1. Proof of Lemma 4.6

*Proof of Lemma 4*.*6*. Let *θ* ∈𝒢_*π*_. For simplicity, write *i* = *π*(1), *j* = *π*(2), and *k* = *π*(3). Since *θ*_*k*_ = 0, Eq. (1.12) implies that

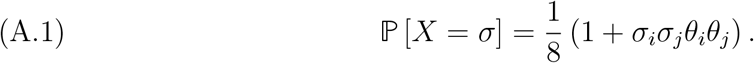

Since the distribution depends only on the product *θ*_*i*_*θ*_*j*_, and not on *θ*_*i*_ or *θ*_*j*_ independently, the log-likelihood restricted to the set *G*_*π*_ may thus be regarded as function of the single variable *x* := *θ*_*i*_*θ*_*j*_ ∈ (0, 1). Plugging Eq. (A.1) into Eq. (1.8), we obtain

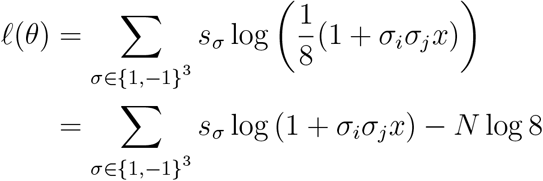

where the second equality follows from ∑_*σ*∈*{*1,−1*}*_ *s*_*σ*_ = *N*. Regrouping terms,

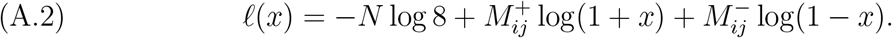

Differentiating gives

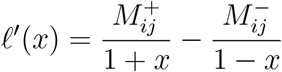

Solving the equation 𝓁^*′*^(*x*) = 0, we find that *f* has at most one critical point on (0, 1), which is the point

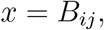

provided that *B*_*ij*_ ∈(0, 1). If this is the case, then *x* is a local maximum by the second derivative test, since

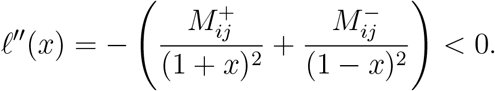

Plugging *x* = *B*_*ij*_ into Eq. (A.2) implies Eq. (4.22), since 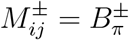 and *B*_*ij*_ = *B*_*π*_.

Eq. (4.23) then follows from Eq. (4.22) along with the observations that 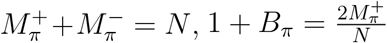, and 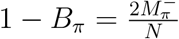.

### A.0.2. Proof of Lemma 4.9

*Proof of Lemma 4*.*9*. We prove the case with *π* = (1), as the proofs for the cases with *π* ∈ *{*(123), (132)*}* are similar. If *ℰ*_int_ = ∅ then there is nothing to show, so henceforth assume *ℰ*_int_ ≠ ∅. By Lemma 4.2, *ℰ*_int_ = *{θ*^∗^*}* and

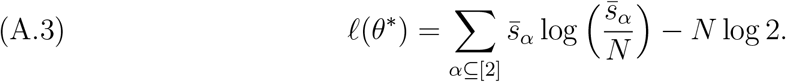

Also by Lemma 4.2 it must be the case that

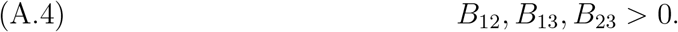

Moreover since *B*_12_ *>* 0, it follows that 𝒢_(1)_ ≠ ∅ by Lemma 4.6.

Let *θ* ∈𝒢_*π*_. By Eq. (4.23),

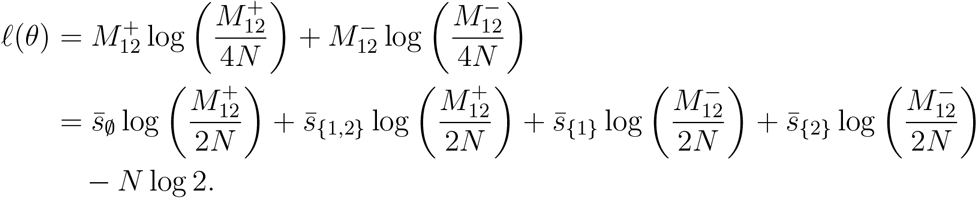

Therefore by Theorem 4.7 and by Eq. (A.3),

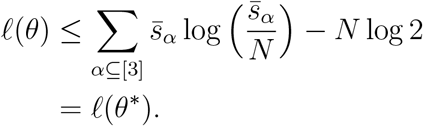

Suppose that equality holds in the above, i.e., that 𝓁(*θ*) = 𝓁(*θ*^∗^). Then by Theorem 4.7,

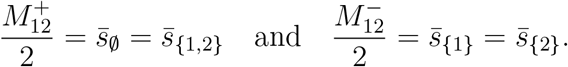

Therefore,

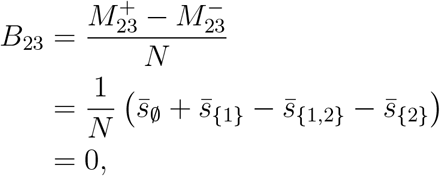

but this contradicts Eq. (A.4). We conclude that 𝓁(*θ*) *<* 𝓁(*θ*^∗^).

### A.0.3. Proof of Lemma 4.12

*Proof*. Since 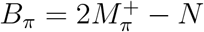 and 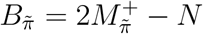, therefore Eq. (4.29) can be rewritten

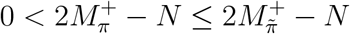

and therefore it holds that

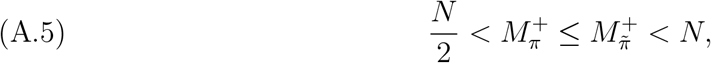

where the last inequality holds by **A.1**.

If 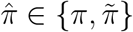 and 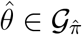, then Eq. (4.23) implies

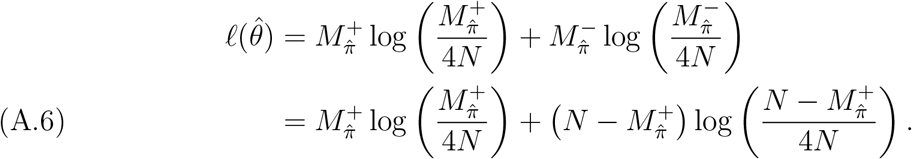

The right-hand side has the form 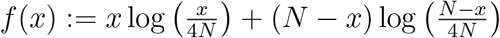. Since

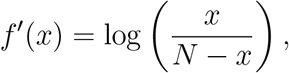

which is positive for all *x* ∈ (*N/*2, *N*), it follows that *f* is strictly increasing on 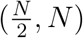. Combining this fact with Eqs. (A.5) and (A.6) implies 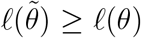, with equality only if 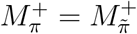, or equivalently 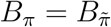.

### A.0.4. Proof of Lemma 4.13

*Proof of Lemma 4*.*13*. If *θ* ∈𝒢_*π′*_ then Lemma 4.6 implies *B*_*π*_ *>* 0. Therefore since 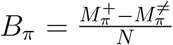 and 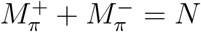, it follows that

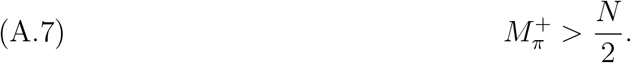

Let 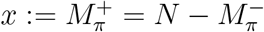. By Eq. (4.23),

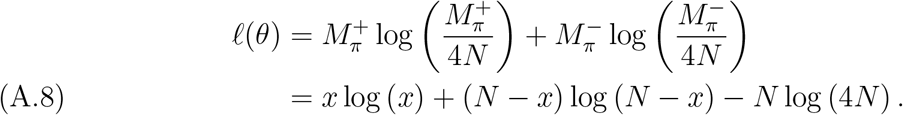

The function *x* ⟼ *x* log (*x*) + (*N* − *x*) log (*N* − *x*) is strictly increasing (on the interval 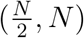, which can be seen by observing that its derivative is 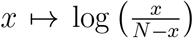, which is positive on this interval. Since 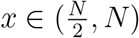 by Eq. (A.7), it follows that

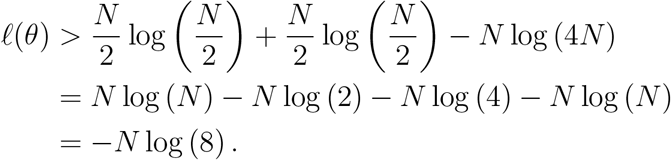

### A.0.5. Proof of Lemma 1.3

*Proof of Lemma 1*.*3*. Let *A* := (*X*_*i*_ : *i* ∈*A*_*e*_) and *B* := (*X*_*i*_ : *i* ∈*A*_*e*_). Write *e* = (*e*_*A*_, *e*_*B*_), and without loss of generality assume that *e*_*A*_ and *e*_*B*_ are labeled such that any path from a leaf in *A*_*e*_ to *e*_*A*_ does not contain *e*_*B*_. Let *Z*_*A*_, *Z*_*B*_ denote the nucleotide states of vertices *e*_*A*_ and *e*_*B*_ respectively. It follows by Eq. (1.2) and symmetry of the process that *Z*_*A*_ and *Z*_*B*_ are independent.

Let 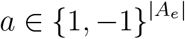 and 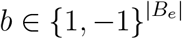. Then using the Markov property,

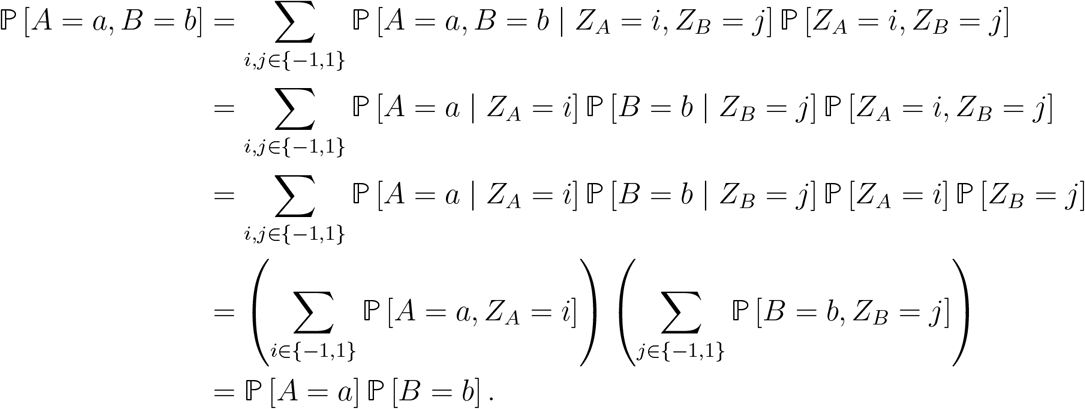

## Appendix B. Code

To compute the characteristic polynomial of *H*_*F*_ in Lemma 4.2, we utilize the following Julia (v1.8.5) code:

~~~
using Homotopy Continuation, LinearAlgebra
@var x y z *λ* c[1:4] t[1:4]
v = [1+ x+y+z,1 -x- y+z,1 - x+y-z, 1+x-y- z] # dummy variable
logL = c’ log .(v) # the log - likelihood (up to a constant)
H = differentiate (differentiate (logL, [x,y, z]),[x,y, z])’
P_char = det(*λ*\*I-H)
P_simplified = expand (subs(P_char, v=>t))
~~~

## Footnotes

1 For trees with more than 3 leaves, a similar interpretation holds using the inequalities of the 4-point condition.

